# Strategy for detecting off-target sites in genome-edited rice

**DOI:** 10.1101/2021.05.28.446070

**Authors:** Jumpei Narushima, Shinya Kimata, Yuh Shiwa, Takahiro Gondo, Satoru Akimoto, Keisuke Soga, Satoko Yoshiba, Kosuke Nakamura, Norihito Shibata, Kazunari Kondo

## Abstract

Genome-editing using the CRISPR-Cas9 system can substantially accelerate crop breeding. Because off-target editing is the main problem with this system, a reliable method for comprehensively detecting off-target sites is required for the editing of food crop genomes. However, a method that accurately predicts off-target sites has not been established. In this study, we performed a SITE-Seq analysis to predict potential off-target sites. SITE-Seq is an unbiased method applicable for the *in vitro* detection of double-strand breaks (DSBs). To analyze SITE-Seq data, we developed a novel Galaxy system, which can perform simple and reproducible analyses without a command line operation. We conducted a SITE-Seq analysis of a rice genome modified by *OsFH15* gRNA-Cas9, and identified 41 DSB sites in the annotated regions. Amplicon-sequencing revealed mutations at one off-target site in the genome-edited rice. The presence of an uncommon protospacer adjacent motif (NTG PAM) likely makes this off-target site difficult to identify using *in silico* methods. Of the six tested programs, only CRISPRdirect predicted this off-target site, but it also predicted 6,080 off-target sites in total. These results suggest the SITE-Seq method presented herein can efficiently predict off-target sites and is useful for assessing the safety of genome-edited food.

## Introduction

Genome-editing technology is a powerful tool used for plant breeding^1^, gene therapy^2^, and drug development^3^. More specifically, the clustered regularly interspaced short palindromic repeats (CRISPR) and CRISPR-associated protein 9 (Cas9) system has been recently applied for improving the traits of various crops, including rice, maize, and tomato^1,4–6^. However, site-directed nucleases, such as Cas9, cleave the target site (on-target) as well as unintended sites (off-target), which is a major concern for the genetic modification of food crops^7–11^. Despite the progress made in genome-editing research, accurately predicting off-target sites with double-strand breaks (DSBs) remains difficult. Comprehensively predicting off-target sites during genome-editing is important for the subsequent assessment of food safety. Although the safety of genome-edited food must be confirmed, there is a lack of a standard protocol for predicting off-target sites.

Off-target sites are currently predicted mainly via *in silico* or experimental methods. The *in silico* methods involve a search for on-target sites based on sequence homology. To date, many *in silico* methods have been developed and are widely used for predicting off-target sites^12–18^. However, it is unclear whether these methods can detect all off-target sites, especially low-homology sites. Furthermore, thresholds for an acceptable number of mismatches and gaps have not been established. In contrast, experimental methods apply next-generation sequencing (NGS) to detect DSBs^7–11,19–26^. These methods can be divided primarily as *in vivo*^19–22^ and *in vitro*^7,8,23–25^ methods. Because *in vivo* methods detect DSBs produced by a ribonucleoprotein (RNP; guide RNA [gRNA]–Cas9 protein complex) in the intracellular environment, predictions are closer to the actual genome-editing situation. However, some off-target sites may be missed because DSBs are repaired in cells^11^. The *in vitro* methods detect DSBs produced by RNP using purified DNA. Although *in vitro* methods are superior to *in vivo* methods in terms of sensitivity and are applicable for diverse species, including animals and plants, the cut sites identified by *in vitro* methods include many false positives^10,11^.

Developing a comprehensive protocol for detecting off-target sites is essential for enhancing the application of genome-editing technology for food production. We believe *in vitro* methods are necessary for identifying off-target sites for genome-edited food because they are unbiased and highly sensitive. In this study, we used the selective enrichment and identification of tagged genomic DNA ends by sequencing (SITE-Seq)^7^, which is an unbiased *in vitro* method, to predict possible off-target cleavages using gRNA designed for rice (*Oryza sativa* L.) as a model crop. Furthermore, we developed a novel system, namely the Galaxy for Cut Site Detection program, which is run on the Docker container and used for SITE-Seq analyses. The results were compared with those obtained from *in silico* methods for predicting off-target sites. Additionally, the utility of SITE-Seq was verified by amplicon-sequencing. A series of analyses revealed one off-target site in genome-edited rice that was difficult to predict using *in silico* methods. Our data suggest that SITE-Seq is an effective method for predicting off-target sites in genome-edited food. Therefore, detecting off-target sites by combining *in silico* methods and SITE-Seq may lead to improved evaluations of the safety of genome-edited food.

## Results

The workflow in this study is presented in Figure 1. First, rice genomic DNA (gDNA) extracted from seedlings was digested with RNP under *in vitro* conditions for the subsequent SITE-Seq analysis. The resulting data were analyzed using the Galaxy for Cut Site Detection program we developed for detecting sites cleaved by RNP. The Galaxy system only requires a sequencing data file, a reference genome file, and a genome annotation file. Any species can be analyzed if a reference genome file is available. Next, we used the CRISPR-Cas9 system to generate genome-edited rice, after which we performed amplicon-sequencing using NGS technology to verify the mutation at the cut sites predicted by SITE-Seq. A negative control was necessary for determining whether the mutation resulted from genome-editing, endogenous single nucleotide polymorphisms, or PCR and/or NGS errors.

**Figure 1.**
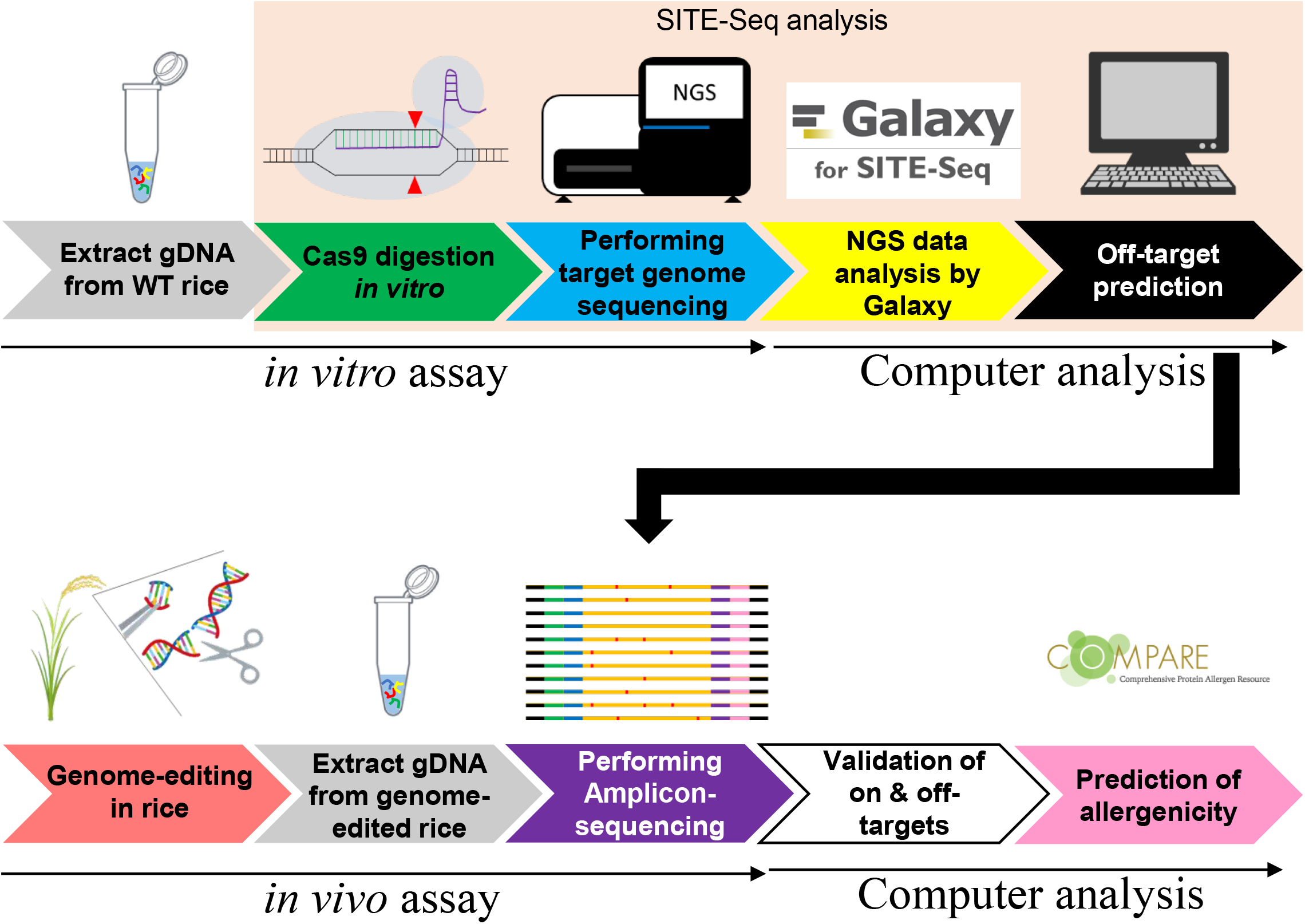
Study workflow

### SITE-Seq analysis on the Galaxy system

Galaxy (https://usegalaxy.org/) is a web-based platform that enables NGS analyses with a graphical user interface^27^. We developed a shell script for the SITE-Seq analysis and combined it with a previously developed program for detecting cut sites^7^. We then extended the pipeline to the Docker-Galaxy system (Fig. 2). The pipeline was as follows: (1) SITE-Seq read data were aligned to the reference genome using Bowtie2, and the data were converted to a bam format using Samtools. (2) The aligned reads in the bam format and the reference genome file were analyzed to detect cut sites using the reported program^7^, and the data were exported in a bed format. (3) Details regarding the cut sites were compared with the information in the reference annotation file, after which the cut sites in annotated genes were extracted using Bedtools. (4) The extracted cut sites were refined using the awk command. Finally, the information was exported as a text file. The cut sites detected via the Galaxy for Cut Site Detection program are listed in the Supplementary Dataset.

**Figure 2.**
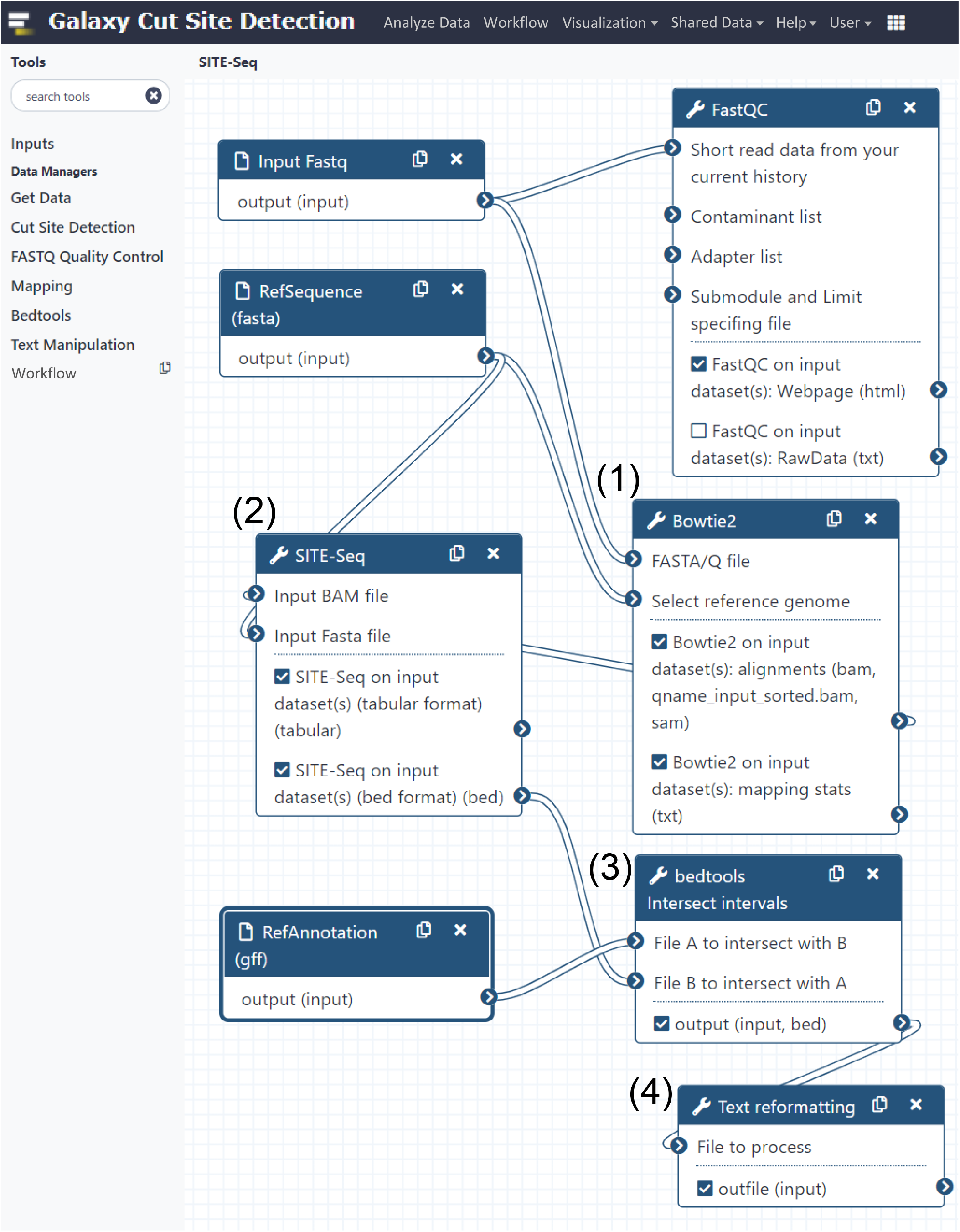
Galaxy for Cut Site Detection pipline.

### Cut sites detected by SITE-Seq

For the SITE-Seq analysis, we focused on the *OsFH15* gene. An earlier study proved that knocking out this gene via CRISPR-Cas9 genome-editing results in decreased rice grain size^28^. In this study, the reported gRNA was used as the basis for editing the rice genome. The DNA libraries for the SITE-Seq analysis were prepared using rice (cv. Nipponbare) gDNA treated with three Cas9 concentrations (64, 256, and 1,024 nM). A total of 219, 484, and 425 cut sites in the whole genome were detected following the 64, 256, and 1,024 nM Cas9 treatments, respectively (Fig. 3a, Supplementary Dataset). These results suggested that the cut sites were comprehensively detected following the treatments with 256 and 1,024 nM Cas9. The annotated regions comprised 41 cut sites. More precisely, 4 sites (two in the exon and two in the untranslated region [UTR]), 12 sites (eight in the exon, two in the intron, and two in the UTR), and 35 sites (20 in the exon, 4 in the intron, and 11 in the UTR) were detected after the 64, 256, and 1,024 nM Cas9 treatments, respectively (Fig. 3b). Regarding these cut sites in the annotated regions, we further investigated nine sites detected at more than one Cas9 concentration. The sites included one on-target and eight off-target sites (Fig. 3c, d and Supplementary Table S1).

**Figure 3.**
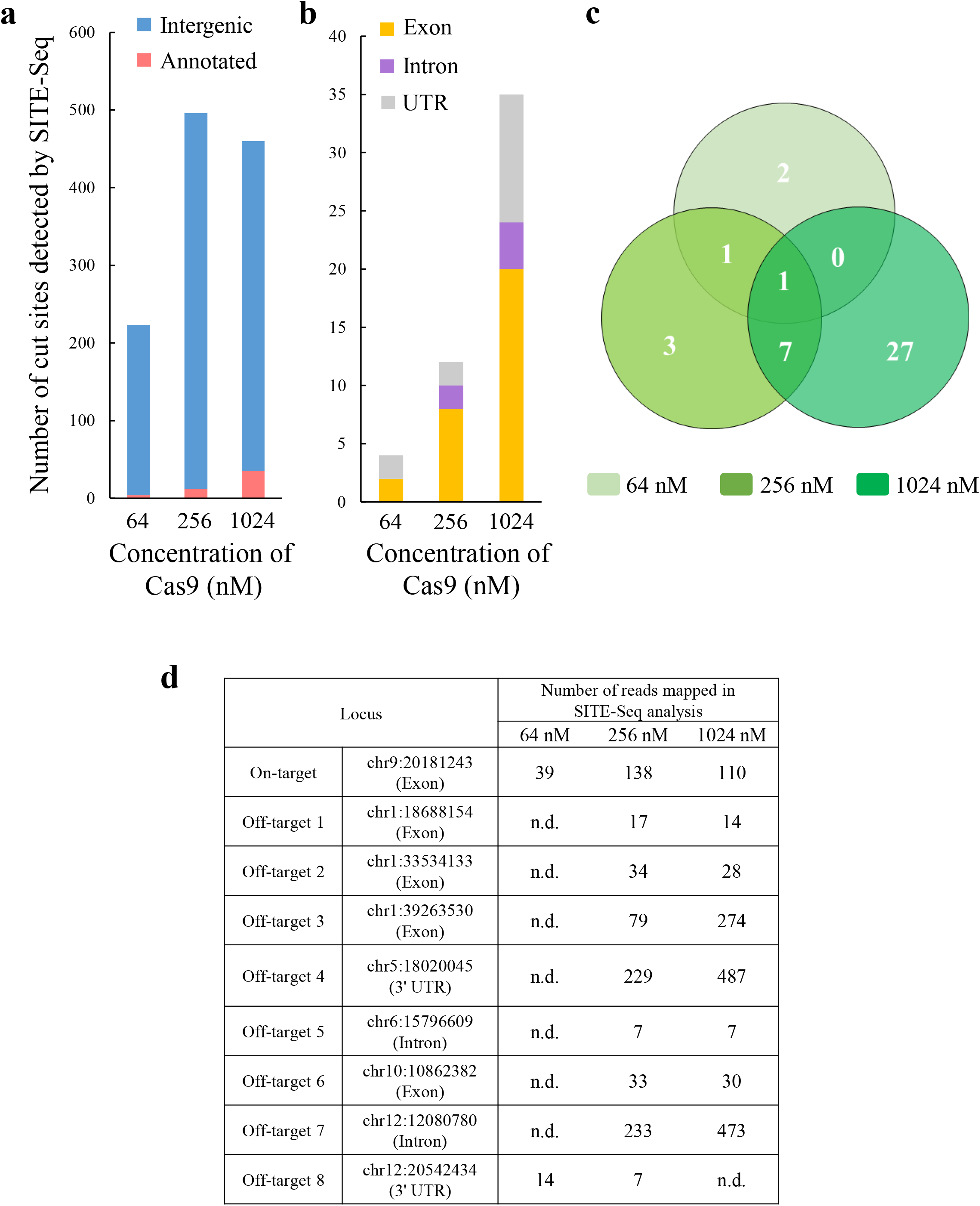
Detection of cut sites by SITE-Seq. (**a**) Bar plot of the number of cut sites detected by SITE-Seq. (**b**) Bar plot of the number of cut sites in (**a**) that are in annotated genome regions. (**c**) Comparison of the number of cut sites in annotated regions detected at various Cas9 concentrations. The on-target site was detected at all Cas9 concentrations. (**d**) Nine cut sites detected at multiple Cas9 concentrations; n.d., not detected.

To determine whether the nine sites predicted by SITE-Seq were digested by RNP, we performed a quantitative PCR (qPCR) analysis of one on-target and three off-target sites (3, 4, and 7) with many mapped reads (Fig. 3d). The qPCR data indicated the Cq values were higher for the digested samples than for the negative control (Supplementary Fig. S1), implying the template DNA quantity decreased because of the digestion by RNP. Next, we used the ΔCq value to calculate the efficiency of the RNP cleavage. At all Cas9 concentrations, the on-target site was efficiently digested (>90%) (Table 1). Off-target sites 3, 4, and 7 were significantly digested (>50%) at 1,024 nM Cas9 (Table 1). Accordingly, the sites predicted by SITE-Seq might be edited by Cas9 under *in vivo* conditions.

**Table 1.** RNP cleavage efficiency for on-target and off-target sites.

| Target | On-target |  |  |  | Off-target 3 |  |  |  | Off-target 4 |  |  |  | Off-target 7 |  |  |  |
| --- | --- | --- | --- | --- | --- | --- | --- | --- | --- | --- | --- | --- | --- | --- | --- | --- |
| Sample | Cas9 concentration (nM) |  |  |  | Cas9 concentration (nM) |  |  |  | Cas9 concentration (nM) |  |  |  | Cas9 concentration (nM) |  |  |  |
|  | NC | 64 | 256 | 1,024 | NC | 64 | 256 | 1,024 | NC | 64 | 256 | 1,024 | NC | 64 | 256 | 1,024 |
| Mean C<sub>q</sub> | 21.85 | 28.33 | 28.34 | 28.74 | 22.06 | 22.09 | 22.40 | 23.21 | 23.23 | 24.34 | 27.76 | 27.85 | 21.82 | 22.10 | 23.46 | 25.53 |
| SD | 0.07 | 0.18 | 0.25 | 0.09 | 0.01 | 0.02 | 0.17 | 0.07 | 0.20 | 0.09 | 0.05 | 0.08 | 0.01 | 0.04 | 0.14 | 0.01 |
| $\Delta C_q$ | – | 6.48 | 6.49 | 6.89 | – | 0.04 | 0.34 | 1.16 | – | 1.12 | 4.53 | 4.63 | – | 0.29 | 1.64 | 3.71 |
| Cleavage efficiency (%) | – | 98.88 | 98.89 | 99.16 | – | 1.96 | 21.26 | 54.95 | – | 53.70 | 95.67 | 95.93 | – | 17.36 | 67.95 | 92.36 |

The C<sub>q</sub> values are described in Supplementary Fig. S1. Cleavage efficiency (%) was calculated using $\Delta C_q$ values.

### Comparison between SITE-Seq and *in silico* programs for predicting off-target sites

We performed an *in silico* analysis to predict off-target sites for *OsFH15*-gRNA. Moreover, we checked whether the nine cut sites detected by SITE-Seq could be predicted by the *in silico* analysis (Table 2). The tested programs were unable to predict off-target sites 1, 2, 5, 6, and 8, probably because of the low homology with the on-target protospacer sequence. Off-target site 3, which contains the NAG protospacer adjacent motif (PAM), was predicted by three programs (Cas-OFFinder^13^, CRISPRdirect^16^, and CRISPR-DT^17^), whereas off-target site 4 was predicted by two programs (Cas-OFFinder and CRISPRdirect). Off-target site 7, which contains the NTG PAM sequence, was predicted by only CRISPRdirect because this program allows mismatches in the PAM sequence. However, 6,080 candidate off-target sites were detected by CRISPRdirect, making it difficult to thoroughly analyze all of these sites (Supplementary Table S2).

**Table 2.**
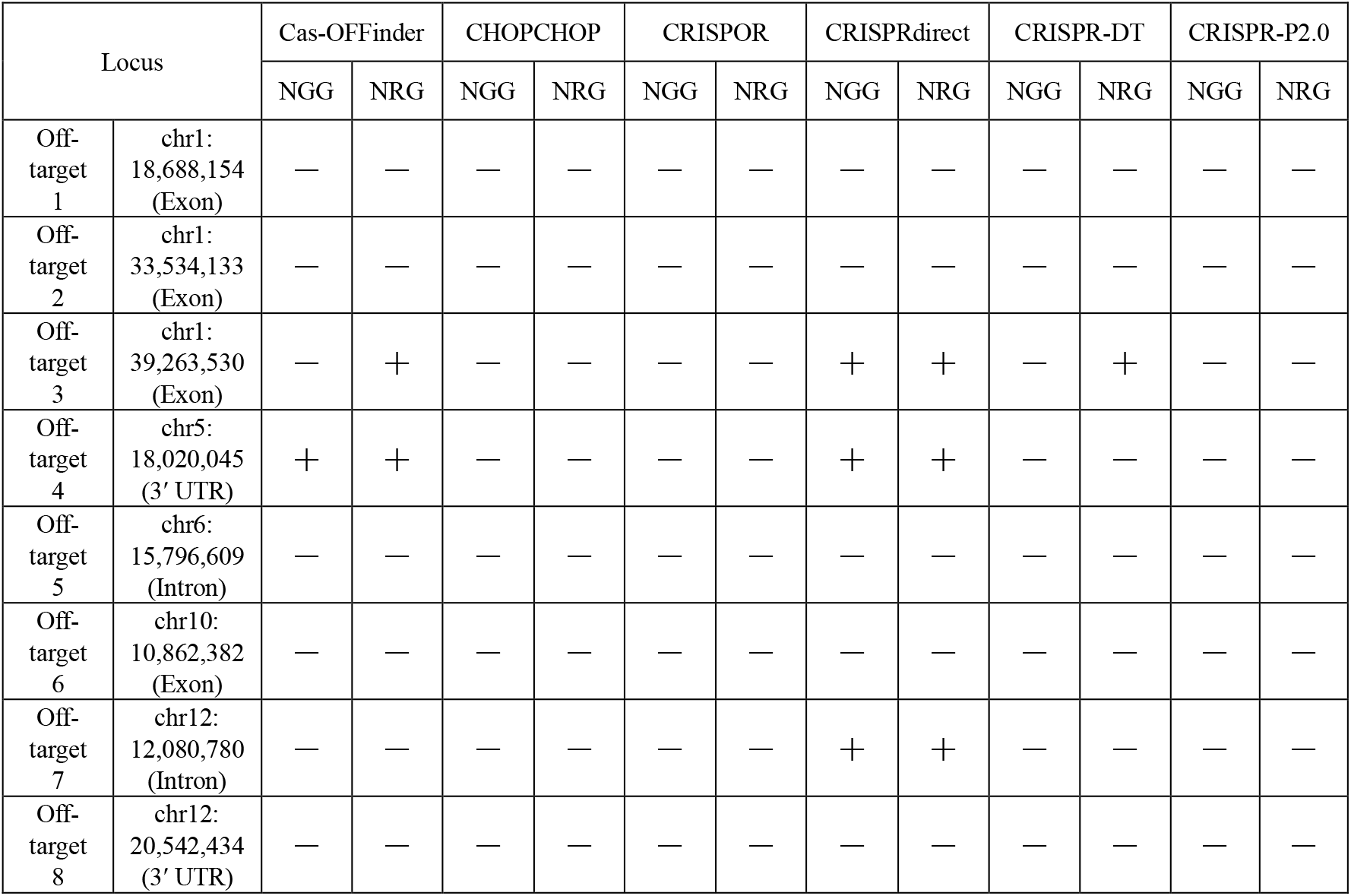
Comparison between SITE-Seq and *in silico* programs for predicting off-target sites

### Validation of potential off-target sites in genome-edited rice calli

To evaluate whether the cut sites detected by SITE-Seq would be digested by Cas9 under *in vivo* conditions, we produced genome-edited rice using *OsFH15*-gRNA. We introduced a plant expression cassette with Cas9 and gRNA into rice calli according to an *Agrobacterium*-mediated transformation method. Transformed calli were selected on the basis of their resistance to hygromycin B, after which they were cultured for 1 or 2 months without any regeneration. First, we performed a T7 endonuclease I (T7E1) assay to detect mutations at one on-target site and three off-target sites (3, 4, and 7). The T7E1 assay is useful for detecting heteroduplexed DNA resulting from mismatches generated by errors while DSBs are being repaired (e.g., by the non-homologous end joining pathway)^29^. The assay of the genome-edited samples revealed T7E1-digested fragments, indicative of genomic modifications, for the on-target site, but not for any of the off-target sites (Supplementary Fig. S2). Because of the low sensitivity of the T7E1 assay, it may not detect low-frequency mutations. Therefore, we sequenced amplicons to more precisely examine the nine cut sites. The amplicon-sequencing results confirmed that the genome-edited samples had various mutations, such as substitutions, deletions, or insertions, at the on-target site. Up to 10.57% of the wild-type reads were in the genome-edited samples cultured for 1 month, whereas more than 95% of the wild-type reads were in the negative control (Fig. 4a, Supplementary Fig. S3). The percentage of wild-type reads in the genome-edited samples cultured for 2 months was less than 0.1% (Supplementary Fig. S4), indicating that the on-target site was almost completely modified under *in vivo* conditions. We then analyzed the eight candidate off-target sites in detail. Of three genome-edited lines cultured for 1 month, we detected a 5-nt deletion at off-target site 7 in only one line (GE1-3); the mutation was absent in the negative control sample (Fig. 4b, Supplementary Fig. S17). Regarding the other candidate off-target sites, no specific mutations were identified in the genome-edited samples (Supplementary Figures S5–S16, S19, and S20). Additionally, specific mutations were undetectable at off-target site 7 in the genome-edited samples cultured for 2 months (Supplementary Fig. S18).

**Figure 4.**
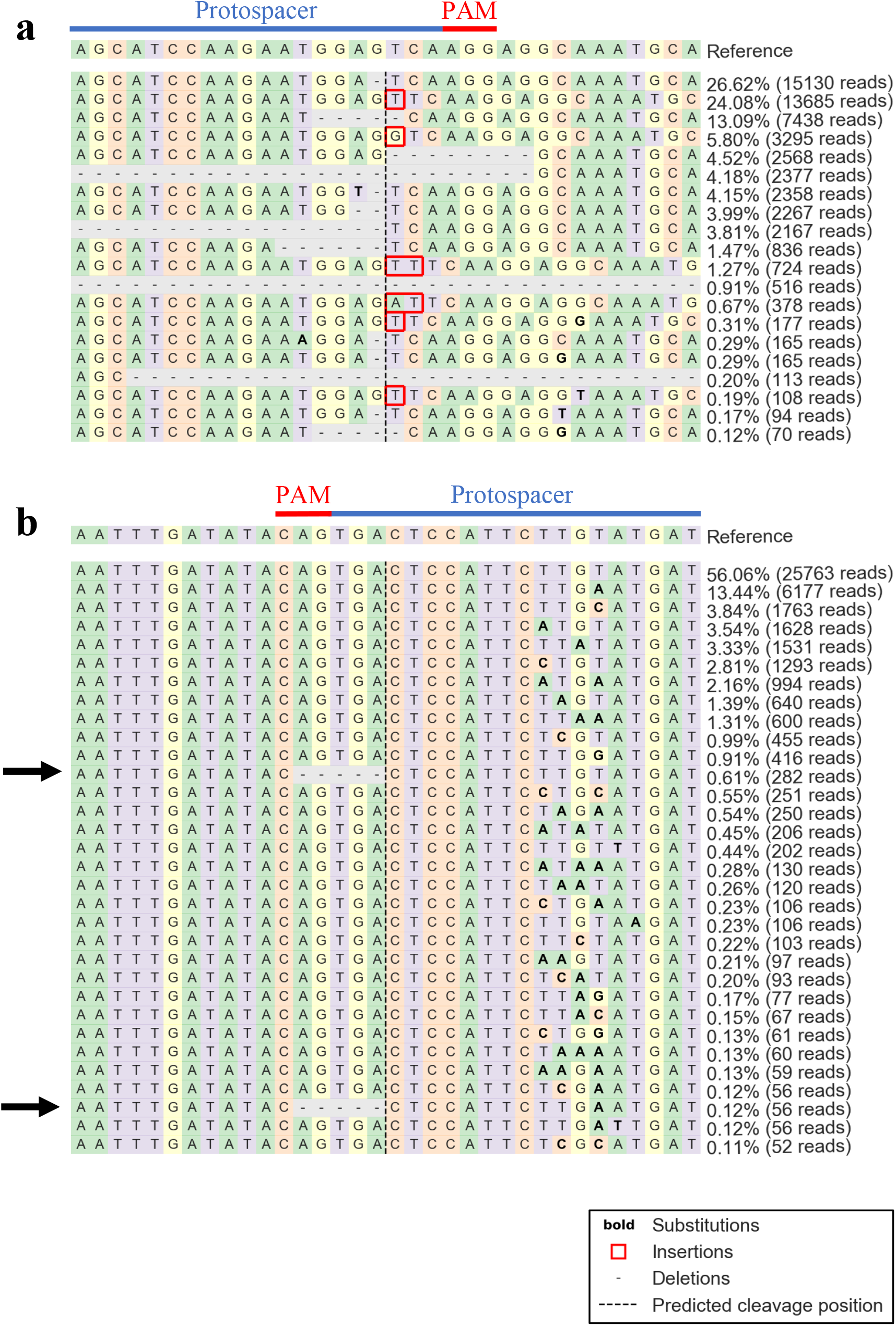
Amplicon-sequencing analysis of the cut sites detected by SITE-Seq. Amplicons were sequenced for the genome-edited and negative control rice calli cultivated for 1 month (n = 3). Regarding the genome-edited samples, the on-target site (**a**) and off-target site 7 (**b**), which had >0.1% mutations, are presented. The read counts and rates are provided for each sequence. The presented data are from the same line (GE1-3). Arrows indicate a 5-nt deletion, which was not detected in the negative control (see Supplementary Fig. S17).

### Assessment of the allergenicity of modified sequences

The gRNA used in this study corresponded to exon 2 of *OsFH15*, whereas off-target site 7 was located in the intron of Os12g0403800. Therefore, we examined the allergenicity of the sequences with a modified on-target site revealed by amplicon-sequencing using the COMPARE Database. According to the 80-mer Sliding Window FASTA Search and 8-mer FASTA Search analyses, none of the tested sequences were allergenic.

## Discussion

In this study, we conducted SITE-Seq and amplicon-sequencing analyses to investigate candidate off-target sites in RNP-digested rice genome and in genome-edited rice calli, respectively. Although the gRNA used in this study was highly specific to the on-target site, a mutation was also detected at off-target site 7, which contains the NTG PAM, under *in vivo* conditions. The digestion of a target site by RNP depends on the recognition of a PAM sequence^30–32^. Earlier studies revealed SpCas9 only weakly modifies genome sequences at sites containing NTG PAM^25,32^, but is more active at sites containing NGG or NAG PAM^25,30–32^. Thus, NTG PAM was considered relatively unimportant for predicting off-target sites. However, our data suggest that a site containing NTG PAM should also be considered as a potential site digested by SpCas9 when assessing the safety of genome-edited food and human therapeutics. Additionally, SITE-Seq is an effective method for evaluating safety because it is an unbiased method that can predict potential cut sites regardless of the PAM sequence. Therefore, in addition to *in silico* analyses, off-target sites should also be predicted experimentally (e.g., SITE-Seq analysis).

We also developed the Galaxy for Cut Site Detection program, which enables highly reproducible analyses using any operating system. Knowledge of command line tools is unnecessary because the system operates on a web browser. Using this pipeline, cut sites in the whole genome or only in annotated regions, including exons, introns, and UTRs, can be detected. Mutations in introns reportedly lead to alternative splicing, such as exon skipping, intron retention, and alternative selection of 5′ or 3′ splice sites, resulting in the production of abnormal proteins^33–35^. Additionally, UTRs are important for translation, localization, and mRNA stability^36,37^. Thus, mutations in UTRs may cause various cellular abnormalities. Therefore, we propose that in addition to exons, the introns and UTRs should also be examined for the presence of off-target sites. The Galaxy for Cut Site Detection program is available online (https://hub.docker.com/repository/docker/nihsdnfi/galaxy-cutsite-detection).

We performed a SITE-Seq analysis to detect candidate sites for off-target DSBs induced by CRISPR-Cas9. Moreover, an amplicon-sequencing analysis was completed to confirm the candidate site mutations. Our results suggest that this strategy can effectively and efficiently identify off-target sites in genome-edited food. To address the possibility the mutations in the edited genome have biological effects, we used an allergen database to investigate whether the modified sequences revealed by amplicon-sequencing are allergenic. A similar evaluation of safety is necessary during the development of genome-edited food. We consider that the procedure developed in the current study (i.e., off-target prediction via SITE-Seq and *in silico* methods, amplicon sequencing, and allergen prediction) will be useful for determining the safety of genome-edited food. However, this approach will need to be refined to increase the accuracy of the predictions.

In addition to CRISPR-Cas9, base editors are believed to be another important technique for genome engineering. Because the currently used base editors contain a catalytically impaired Cas9 fused to a single-stranded DNA deaminase enzyme, they induce targeted point mutations without DSBs. Hence, the generation of insertions and deletions at off-target sites by base editors is likely suppressed^38^, whereas single-nucleotide variants are generated throughout the genome^39,40^. On the basis of Digenome-seq^23^, experimental methods for detecting off-target genome-editing have been developed for cytosine and adenine base editors^41,42^. Because these methods involving Cas9 nickase and endonuclease V or VIII produce blunt-ended DSBs as an intermediate, SITE-Seq may be a useful addition to these off-target detection methods designed for base editors. Furthermore, compared with methods based on Digenome-seq, SITE-Seq may detect off-target sites with a lower sequence coverage because it enriches DSBs using biotinylated oligonucleotides. In future studies, we will improve the Galaxy for Cut Site Detection program so that it can detect off-target sites resulting from base editing.

## Methods

### Plant materials and callus induction

Rice (*Oryza sativa* L. ssp. *japonica* cv. Nipponbare; JP: 222430) seeds were provided by the Genetic Resources Center of the National Agriculture and Food Research Organization in Japan. To extract DNA from the wild-type rice used for the SITE-Seq analysis, surface-sterilized seeds were germinated on filter paper moistened with sterile distilled water, after which they were transferred to Murashige and Skoog medium^43^ supplemented with myoinositol (10 mg/L), glycine (200 μg/L), nicotinic acid (50 μg/L), pyridoxine hydrochloride (50 μg/L), thiamine hydrochloride (10 μg/L), pH 5.8, and 3% sucrose. Seedlings were incubated at 28 °C with an 8-h light/16-h dark photoperiod. To generate rice calli, seeds were surface-sterilized and placed on N6 medium^44^ supplemented with 2,4-dichlorophenoxyacetic acid (2 mg/L), proline (10 mM), casein hydrolysate (300 mg/L), sucrose (30 g/L), and Gelrite (3 g/L) (N6D medium). Calli were induced for 3 weeks. Small, vigorously dividing calli were subcultured at 14-day intervals. The calli were incubated at 28 °C in darkness and then transformed.

### Construction of the CRISPR-Cas9-related vector and *Agrobacterium*-mediated transformation

A vector based on pRGEB32 (Addgene, Watertown, MA, USA, plasmid #63142)^45^ was used for genome-editing. The pair of oligonucleotides used for introducing the gRNA cassette into the vector is listed in Supplementary Table S3. The constructed vector was transferred into *A. tumefaciens* strain EHA105 cells, which were then used to transform rice calli as described by Ozawa^46^. Negative control calli were transformed with the empty pRGEB32 vector (i.e., without the gRNA cassette). After a 21-day infection, transformed calli were selected on medium containing 50 mg/L hygromycin B. Independent lines were subcultured on fresh medium at 14-day intervals. The transformed callus lines were cultured for 1 or 2 months and then used for the genome-editing analysis.

### SITE-Seq analysis

The SITE-Seq analysis was performed using the MiSeq system and MiSeq Reagent Kit (version 3) (150 cycles) (Illumina, Inc., San Diego, CA, USA) to produce 1 × 151-bp single-end reads. Specific details regarding the methods used are provided in the Supplementary Methods. The oligonucleotides used for the SITE-Seq analysis are listed in Supplementary Table S3.

### Galaxy for Cut Site Detection program

Referring to a published report^7^, we wrote a novel shell script to analyze the SITE-Seq data (see Supplementary Script). It includes an additional function that extracts cut sites in annotated regions. Read quality was checked using FastQC (version 0.11.9), after which the reads were aligned to the reference genome using Bowtie2^47^ (version 2-4.2). The sam file for the aligned reads was converted to a bam file before the sorting and indexing step using Samtools^48^ (version 1.12). The sorted bam file and the original python program described in SITE-Seq Supplementary Software^7^ were used for detecting cut sites. The resulting cut site data were converted to a bed file, after which cut sites within annotated regions were called using Bedtools^49^ (version 2.30.0). Furthermore, we built a custom Galaxy for Cut Site Detection program, which runs on a Docker container, from the developed shell script. The IRGSP-1.0 reference genome and genome annotation files were downloaded from RAP-DB (https://rapdb.dna.affrc.go.jp/download/irgsp1.html) in November 2018. The cut sites detected by the SITE-Seq analysis are listed in the Supplementary Dataset.

### Prediction of off-target sites using *in silico* methods

The following programs, which are applicable for analyzing the rice reference genome, were selected for predicting off-target sites: Cas-OFFinder^13^ (http://www.rgenome.net/cas-offinder/), CHOPCHOP^14^ (https://chopchop.cbu.uib.no/), CRISPOR^15^ (http://crispor.tefor.net/crispor.py), CRISPRdirect^16^ (https://crispr.dbcls.jp/), CRISPR-DT^17^ (http://bioinfolab.miamioh.edu/CRISPR-DT/), and CRISPR-P 2.0^18^ (http://crispr.hzau.edu.cn/CRISPR2/). The off-target sites for SpCas9 (both NGG and NRG PAM) were predicted using default parameters.

### qPCR analysis

The qPCR analysis was performed using the LightCycler 480 System II (Roche Diagnostics, Rotkreuz, Switzerland) to evaluate RNP cleavage efficiency. The RNP digestion was completed in a 25-μL reaction mixture containing recombinant SpCas9 (Themo Fisher Scientific, Waltham, MA, USA), gRNA, and 1.5 μg rice gDNA. The final Cas9 concentrations were 64, 256, and 1,024 nM. The method for synthesizing gRNA is described in the Supplementary Methods and details regarding the oligonucleotides used are listed in Supplementary Table S3. The digestion by RNP was performed at 37 °C for 16 h, after which 44 μg RNase A (Nippon Gene Co., Ltd., Tokyo, Japan) was added to the mixture, which was then incubated at 37 °C for 20 min. The reaction was terminated by adding 5 μg protease K (Wako) and incubating the mixture at 55 °C for 20 min. Finally, protease K was deactivated via an incubation at 95 °C for 10 min. For the negative control, water was added instead of gRNA. The qPCR was performed in a 25−μL reaction mixture that included 12.5 μL THUNDERBIRD SYBR qPCR Mix (Toyobo Co., Ltd., Osaka, Japan), 0.5μM each primer, and 5 ng RNP-digested DNA. The qPCR primers are listed in Supplementary Table S3. The ΔCq values were calculated by subtracting the Cq value of the negative control from the Cq value of the genome-edited sample. Cleavage efficiency (*E*) was calculated using the following equation:

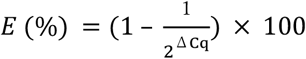

### T7 endonuclease I assay

The on-target and off-target sites detected by SITE-Seq were amplified from genome-edited rice DNA using specific primers (Supplementary Table S3). The PCR products were analyzed by 2% agarose gel electrophoresis and purified using the NucleoSpin Gel and PCR Clean-up kit (Macherey-Nagel, GmbH & Co. KG, Dueren, Germany). Approximately 200 ng DNA products were mixed with NEBuffer 2 (50 mM NaCl, 10 mM Tris-HCl, 10 mM MgCl_2_, and 1 mM dithiothreitol, pH 7.9) and incubated at 95 °C for 5 min before gradually decreasing the temperature to 25 °C (0.1 °C/s). After adding 2.5 U T7 endonuclease I (New England Biolabs Inc., Ipswich, MA, USA), the mixture was incubated at 25 °C for 15 min before 1.5 μL 0.25 M EDTA was added to terminate the reaction. The reaction products were analyzed by 2% agarose gel electrophoresis.

### Amplicon-sequencing

Nine cut sites detected by SITE-Seq were amplified from genome-edited rice callus DNA using specific primers (Supplementary Table S3). The DNA libraries were prepared as described in the Supplementary Methods. Amplicons were sequenced using the iSeq 100 system and the iSeq 100 i1 Reagent (300 Cycles) (Illumina) to generate 2 × 151-bp paired-end reads. The CRISPResso2^50^ program (version 2.0.31; https://github.com/pinellolab/CRISPResso2) was used to analyze the mutation rate greater than 0.1%.

### Allergenicity prediction

The allergenicity of the modified sequences revealed by amplicon-sequencing was predicted using the COMPARE 2020 database released on January 29, 2020 (https://comparedatabase.org/). More specifically, the allergenicity was assessed according to the 80-mer Sliding Window FASTA Search and the 8-mer FASTA Search programs.

## Supporting information

Supplementary Script

## Acknowledgments

This study was supported in part by the Ministry of Health, Labor and Welfare (H30-Syokuhin-Ippan-002 and 21KA1002 to K.K.), and Japan Society for the Promotion of Science and the Ministry of Education, Culture, Sports, Science and Technology (JSPS/MEXT KAKENHI Grants Number JP20K05901 to K.S., JP21K06075 to N.S., and JP21H02469 to K.K.). We thank Edanz (https://jp.edanz.com/ac) for editing a draft of this manuscript.

## Author contributions statement

J.N., S.K., K.N., and K.K. designed the experiments. J.N., S.K., and K.N. performed the experiments. T.G. performed the rice transformations. Y.S. and S.A. developed the Galaxy for Cut Site Detection program. J.N. analyzed the data. J.N., N.S., and K.K. wrote the manuscript. All authors reviewed the manuscript.

## Data availability

All data generated or analyzed during this study are included in this published article and its Supplementary Information files. Nucleotide sequence data are available in the DNA Data Bank of Japan Sequence Read Archive under accession number PRJDB11507.

## Additional information

### Supplementary information

Supplementary Figures S1–S20

Supplementary Tables S1–S3

Supplementary Methods

Supplementary Script

Supplementary Dataset

