## Supplementary Figures and Tables for "Strategy for detecting off-target sites in genome-edited rice"

| Locus | | Cut site sequence predicted by SITE-Seq analysis. |
|---|---|---|
| On-target | chr9: 20181243<br>(Exon) | ----AGCATCCAAGAATGGAGTCA-----------------<br>GGAGAGCATCCAAGAATGGAGTCAAGGAGGCAAATGCAGCT<br> ******************** |
| Off-target 1 | chr1: 18688154<br>(Exon) | -----AGCATCCAAGAATGGAGTCA----------------<br>ATTGTTGTTTCTTAAAGTCTGGTTGGAGTTTCTTAAGTTTC<br> * ** * * * ** |
| Off-target 2 | chr1: 33534133<br>(Exon) | --AGCATCCAAGAATGGAGTCA-------------------<br>ATGCCAGCTCTGACCGAAATCTTTGGAGATGATTCTGTATT<br> ** * ** * * ** |
| Off-target 3 | chr1: 39263530<br>(Exon) | -AGC--ATCCAAGAATGGAGTCA------------------<br>CAACTGAACCAAGAATGGAGCCAGAGAGTGAAGTGCAATAC<br> * * * ************ ** |
| Off-target 4 | chr5: 18020045<br>(3' UTR) | -----AGCATCCAAGAATGGAGTCA------------------<br>ATAGGAGCATC--AGAAGGGAGCCATGGTTGCTTAAGAACATC<br> ****** * ** **** ** |
| Off-target 5 | chr6: 15796609<br>(Intron) | AGCATCCAA--GAAT-GGAGTCA------------------<br>AGAAAAGAAACTAATAGAAGTCGAATACGCGGAATGGTAGT<br>** * ** *** * **** |
| Off-target 6 | chr10: 10862382<br>(Exon) | ---AGCATCCAAGAATGGAGTCA------------------<br>TATGCCAGCTCTGACCGAAATCTTTGGAGATGATTCTGTAT<br> ** * ** * * ** |
| Off-target 7 | chr12: 12080780<br>(Intron) | ---AGCATCCAAGAATGGAGTCA------------------<br>GAAATCATACAAGAATGGAGTCACTGTATATCAAATTGTTA<br> * *** ************** |
| Off-target 8 | chr12: 20542434<br>(3' UTR) | AGCATCCA--AGAATGGAGTCA-------------------<br>AGCTAGCAACAAAGCGGAGTCCCAGGGTCTAGCCCTTCTGA<br>*** ** * * ****** |

**Supplementary Table S1.** List of sequences of cut sites detected by SITE-Seq. Cut site sequences were compared homology with protospacer sequence of on-target site using ClustalW. Green characters indicate predicted PAM. There was no PAM around predicted protospacer sequence in off-target site 5.

| | Cas-OFFinder | | CHOPCHOP | | CRISPOR | | CRISPRdirect (20mer+PAM) | | CRISPRdirect (12mer+PAM) | | CRISPRdirect (8mer+PAM) | | CRISPR-DT | | CRISPR-P v2.0 | |
|---|---|---|---|---|---|---|---|---|---|---|---|---|---|---|---|---|
| | NGG | NRG | NGG | NRG | NGG | NRG | NGG | NRG | NGG | NRG | NGG | NRG | NGG | NRG | NGG | NRG |
| 0 mismatch or gap | 1 | 1 | 0 | 0 | 0 | - | 1 | 1 | 4 | 7 | 720 | 1176 | 1 | 1 | 0 | 0 |
| 1 | 0 | 0 | 0 | 0 | 0 | - | 0 | 0 | 176 | 376 | 33349 | 63563 | 0 | 0 | 0 | 0 |
| 2 | 4 | 5 | 0 | 0 | 0 | - | 0 | 0 | 7755 | 16028 | 721070 | 1357067 | 0 | 0 | 0 | 0 |
| 3 | 6 | 14 | 0 | 0 | 1 | - | 11 | 21 | 143737 | 289409 | - | - | 0 | 2 | 0 | 0 |
| 4 | 112 | 350 | 0 | 0 | 17 | - | 333 | 681 | - | - | - | - | 16 | 33 | 20 | 20 |
| 5 | 2076 | 4439 | 0 | 0 | - | - | 5735 | 11131 | - | - | - | - | 56 | 102 | 0 | 0 |
| Total | 2199 | 4809 | 0 | 0 | 18 | - | 6080 | 11834 | 151672 | 305820 | 755139 | 1421806 | 73 | 138 | 20 | 20 |

**Supplementary Table S2.** Number of off-target candidate sites predicted by computational programs.

**Supplementary Table S3.** List of oligonucleotides used in this study.

| Oligonucleotide name | Sequence (5' - 3') |
|---|---|
| Introducing gRNA casset into pRGEB32 | |
| FH15_gRNA_in_pRGEB32_Fwd | GGCAGCATCCAAGAATGGAGTCA |
| FH15_gRNA_in_pRGEB32_Rev | AAACTGACTCCATTCTTGGATGC |
| gRNA synthesis | |
| OsFH15_gRNA_Fwd | CGATGTAATACGACTCACTATAGGAGCATCCAAGAATGGAGTCAGTTTTAGAGCTATGCTGAAA |
| gRNA_Aend | AAGCACCGACTCGGTGCCACTTTTTCAAGTTGATAACGGACTAGCCTTATTTTAACTTGCTATGCTTTTCAGCATAGCTCTAAAACT |
| DNA Library prep for SITE-Seq | |
| Adapter1_Fwd | [BioON]GTTGACATGCTGGATTGAGACTTCCTACACTCTTTCCCTACACGACGCTCTTCCGATCT |
| Adapter1_Rev | GATCGGAAGAGCGTCGTGTAGGGAAAGAGTGTAGGAAGTCTCAATCCAGCATGTCAAC |
| Adapter2_N7_Fwd | [PHO]GTCGTATTAGTAGTANNNNNNNAGATCGGAAGAGCACACGTCTGAACTCC |
| Adapter2_N6_Fwd | [PHO]GTCGTATTAGTAGTANNNNNNAGATCGGAAGAGCACACGTCTGAACTCC |
| Adapter2_N5_Fwd | [PHO]GTCGTATTAGTAGTANNNNNAGATCGGAAGAGCACACGTCTGAACTCC |
| Adapter2_Rev | ACTACTAATACGACT |
| Recovery_PCR_Fwd | GGAGTTCAGACGTGTGCTC |
| Recovery_PCR_Rev | GTTGACATGCTGGATTGAGACTTC |
| DNA Library prep for amplicon-sequencing | |
| Amp_Adapter1_On_Fwd | CACTCTTTCCCTACACGACGCTCTTCCGATCTTCGCGATGTTCTGCGATTTC |
| Amp_Adapter1_On_Rev | GGAGTTCAGACGTGTGCTCTTCCGATCTGAGCTCCCCACGCGATATC |
| Amp_Adapter1_Off1_Fwd | CACTCTTTCCCTACACGACGCTCTTCCGATCTGACATATTCCTCCATTAACACAAAAG |
| Amp_Adapter1_Off1_Rev | GGAGTTCAGACGTGTGCTCTTCCGATCTTAGAGATCCATTGGGGCAGGC |
| Amp_Adapter1_Off2_Fwd | CACTCTTTCCCTACACGACGCTCTTCCGATCTCTAAAGTTCCTCCACCAAATTGC |
| Amp_Adapter1_Off2_Rev | GGAGTTCAGACGTGTGCTCTTCCGATCTCATATCCACGCTGGTACAGTAG |
| Amp_Adapter1_Off3_Fwd | CACTCTTTCCCTACACGACGCTCTTCCGATCTCACCAGTGGATTTGTGGAGG |
| Amp_Adapter1_Off3_Rev | GGAGTTCAGACGTGTGCTCTTCCGATCTGCCATCCAGCTTCTTGAGTAAC |
| Amp_Adapter1_Off4_Fwd | CACTCTTTCCCTACACGACGCTCTTCCGATCTGGGCATGATAAGTCTGTTCCTC |
| Amp_Adapter1_Off4_Rev | GGAGTTCAGACGTGTGCTCTTCCGATCTCAGCAGCAGAGCCACAGAC |
| Amp_Adapter1_Off5_Fwd | CACTCTTTCCCTACACGACGCTCTTCCGATCTGCAAAATTAGCACAGAGACAACC |
| Amp_Adapter1_Off5_Rev | GGAGTTCAGACGTGTGCTCTTCCGATCTCGAGTTAATTTGCCTTGAGACTATC |
| Amp_Adapter1_Off6_Fwd | CACTCTTTCCCTACACGACGCTCTTCCGATCTAGGTGCATTACCCCAAGGATG |
| Amp_Adapter1_Off6_Rev | GGAGTTCAGACGTGTGCTCTTCCGATCTAGGTAAGTTAGAAGGGGAACGC |
| Amp_Adapter1_Off7_Fwd | CACTCTTTCCCTACACGACGCTCTTCCGATCTGGGTGGATTGGTTTGTATGC |
| Amp_Adapter1_Off7_Rev | GGAGTTCAGACGTGTGCTCTTCCGATCTATCCAATTCCAAACTATAAACCGC |
| Amp_Adapter1_Off8_Fwd | CACTCTTTCCCTACACGACGCTCTTCCGATCTGTTCAAAGCCCTAACCAAGTTC |
| Amp_Adapter1_Off8_Rev | GGAGTTCAGACGTGTGCTCTTCCGATCTTTCGCAACCAGTTAGACCTC |
| Index primers | |
| Index-UDI0001_Fwd | AATGATACGGCGACCACCGAGATCTACACAGCGCTAGACACTCTTTCCCTACACGACG |
| Index-UDI0001_Rev | CAAGCAGAAGACGGCATACGAGATAACCGCGGGTGACTGGAGTTCAGACGTGTGCTC |
| Index-UDI0002_Fwd | AATGATACGGCGACCACCGAGATCTACACGATATCGAACACTCTTTCCCTACACGACG |
| Index-UDI0002-Rev | CAAGCAGAAGACGGCATACGAGATGGTTATAAGTGACTGGAGTTCAGACGTGTGCTC |
| Index-UDI0003_Fwd | AATGATACGGCGACCACCGAGATCTACACCGCAGACGACACTCTTTCCCTACACGACG |
| Index-UDI0003-Rev | CAAGCAGAAGACGGCATACGAGATCCAAGTCCGTGACTGGAGTTCAGACGTGTGCTC |
| Index-UDI0004_Fwd | AATGATACGGCGACCACCGAGATCTACACTATGAGTAACACTCTTTCCCTACACGACG |
| Index-UDI0004-Rev | CAAGCAGAAGACGGCATACGAGATTTGGACTTGTGACTGGAGTTCAGACGTGTGCTC |
| Index-UDI0005_Fwd | AATGATACGGCGACCACCGAGATCTACACAGGTGCGTACACTCTTTCCCTACACGACG |
| Index-UDI0005-Rev | CAAGCAGAAGACGGCATACGAGATCAGTGGATGTGACTGGAGTTCAGACGTGTGCTC |
| Index-UDI0006_Fwd | AATGATACGGCGACCACCGAGATCTACACGAACATACACACTCTTTCCCTACACGACG |
| Index-UDI0006-Rev | CAAGCAGAAGACGGCATACGAGATTGACAAGCGTGACTGGAGTTCAGACGTGTGCTC |
| Index-UDI0007_Fwd | AATGATACGGCGACCACCGAGATCTACACACATAGCGACACTCTTTCCCTACACGACG |
| Index-UDI0007-Rev | CAAGCAGAAGACGGCATACGAGATCTAGCTTGGTGACTGGAGTTCAGACGTGTGCTC |
| Index-UDI0008_Fwd | AATGATACGGCGACCACCGAGATCTACACGTGCGATAACACTCTTTCCCTACACGACG |
| Index-UDI0008-Rev | CAAGCAGAAGACGGCATACGAGATTCGATCCAGTGACTGGAGTTCAGACGTGTGCTC |
| Index-UDI0009_Fwd | AATGATACGGCGACCACCGAGATCTACACCCAACAGAACACTCTTTCCCTACACGACG |
| Index-UDI0009-Rev | CAAGCAGAAGACGGCATACGAGATCCTGAACTGTGACTGGAGTTCAGACGTGTGCTC |
| Index-UDI00010-Fwd | AATGATACGGCGACCACCGAGATCTACACTTGGTGAGACACTCTTTCCCTACACGACG |
| Index-UDI00011-Fwd | AATGATACGGCGACCACCGAGATCTACACCGCGGTTCACACTCTTTCCCTACACGACG |
| Index-UDI00012-Fwd | AATGATACGGCGACCACCGAGATCTACACTATAACCTACACTCTTTCCCTACACGACG |
| qPCR | |
| FH15_On_Fwd | AGCATCCAAGAATGGAGTCAAG |
| FH15_On_Rev | TCAGTTCCAAGGCTCTCTGC |
| FH15_Off3_Fwd | ATTGCACTTCACTCTCTGGCTC |
| FH15_Off3_Rev | CTTGTGCTGTGAGAAGTGTGC |
| FH15_Off4_Fwd | ATGTTCTTAAGCAACCATGGCTC |
| FH15_Off4_Rev | CAGCGCTGAAACTGGCTC |
| FH15_Off7_Fwd | TGCTAACAATTTGATATACAGTGACTC |
| FH15_Off7_Rev | TCAGATCGGTGTATCCAATTCC |
| T7E1 assay | |
| FH15_T7E1_On_Fwd | CTTGGTGATTGGATTTAGCAGCAG |
| FH15_T7E1_On_Rev | CATCCAGGAGAGCCTTGCAG |
| FH15_T7E1_Off3_Fwd | TCCTCCTCACCAAATTCGGTAAC |
| FH15_Off3_Rev | CTTGTGCTGTGAGAAGTGTGC |
| FH15_T7E1_Off4_Fwd | TCGCGGAGAATTACCAGCAG |
| FH15_Off4_Rev | CAGCGCTGAAACTGGCTC |
| FH15_T7E1_Off7_Fwd | GCACAGGAGAGCATTAGAGTG |
| FH15_Off7_Rev | TCAGATCGGTGTATCCAATTCC |

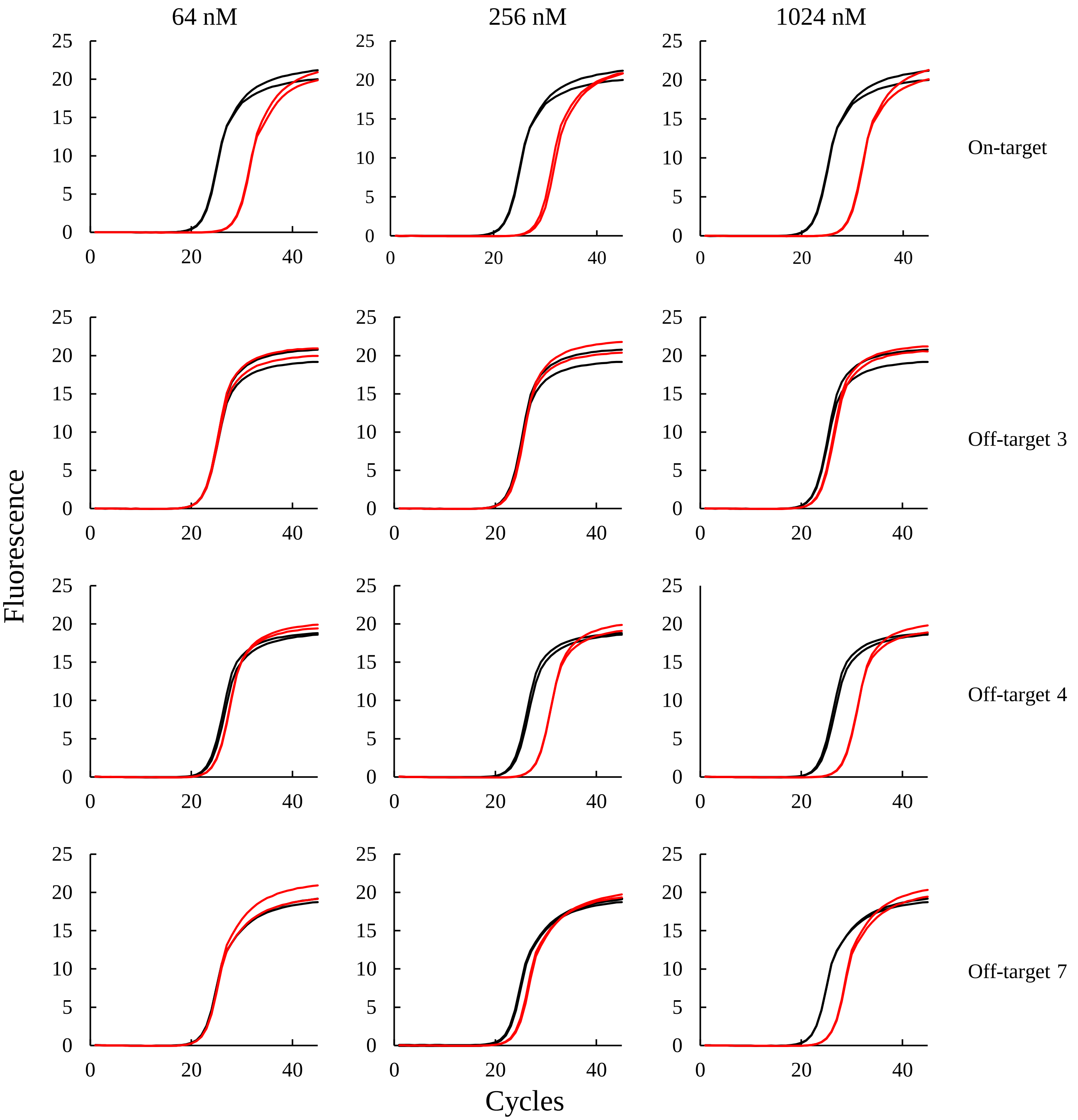


**Supplementary Figure S1.** qPCR assay to quantify RNP cleavage efficiency. Top panels shows analysis of on-target site digested by 64, 256, 1024 nM RNP, respectively. Other panels show qPCR analysis for off-target sites detected by SITE-Seq. The DNA templates were digested by only Cas9 (negative control, black) or gRNA and Cas9 (digested sample,  red) at 37°C for 16 h *in vitro* condition.

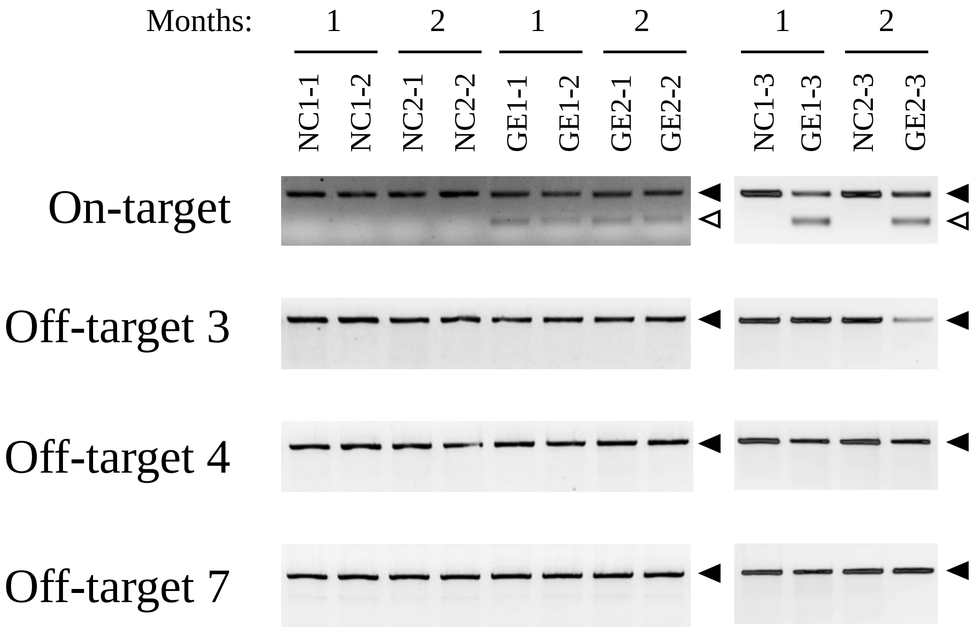


**Supplementary Figure S2.** T7E1 assay using gDNA extracted from genome-edited rice callus cultivated for one or two months (n=3). NC (negative control) were introduced genome-edited vector which contained Cas9 but not gRNA cassette. GE (genome-edited sample) were introduced genome-edited vector which contained gRNA and Cas9. Filled arrowheads indicate undigested bands. Open arrowheads indicate digested bands.

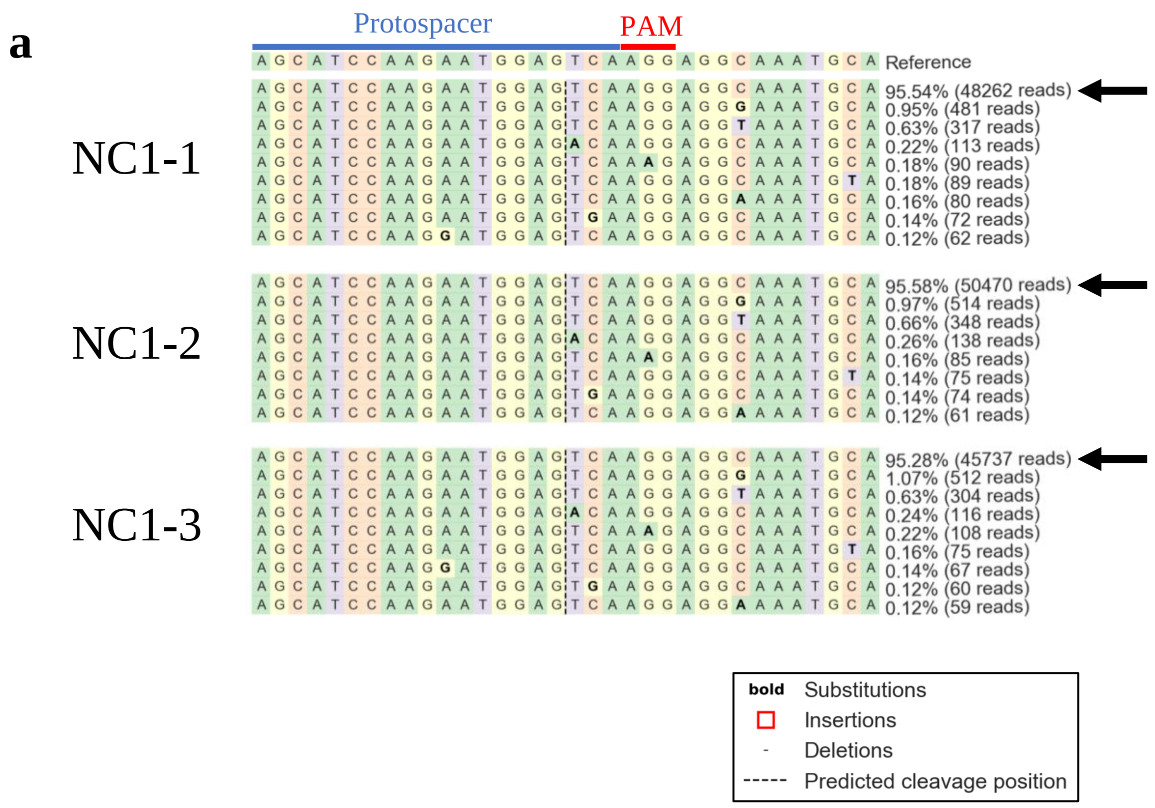


**Supplementary Figure S3**. Amplicon-sequencing analysis for on-target site in negative control (**a**) and genome-edited (**b**) rice callus cultivated for one month (n=3). The site that accumulated > 0.1% mutations are shown. The reads counts and its rate were displayed on each sequences. Arrows represent reference sequence.

**Supplementary Figure S3**. (continued)

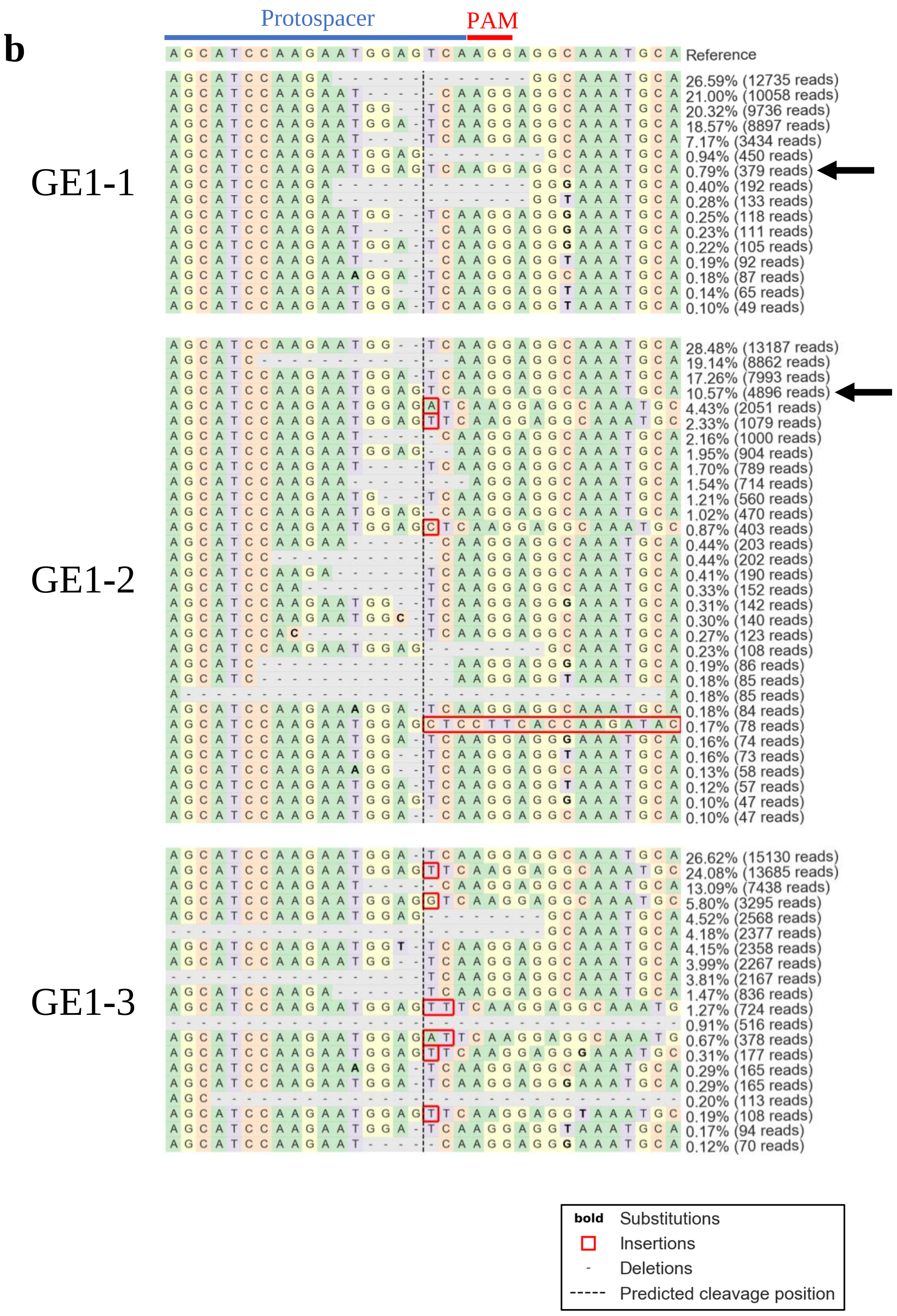

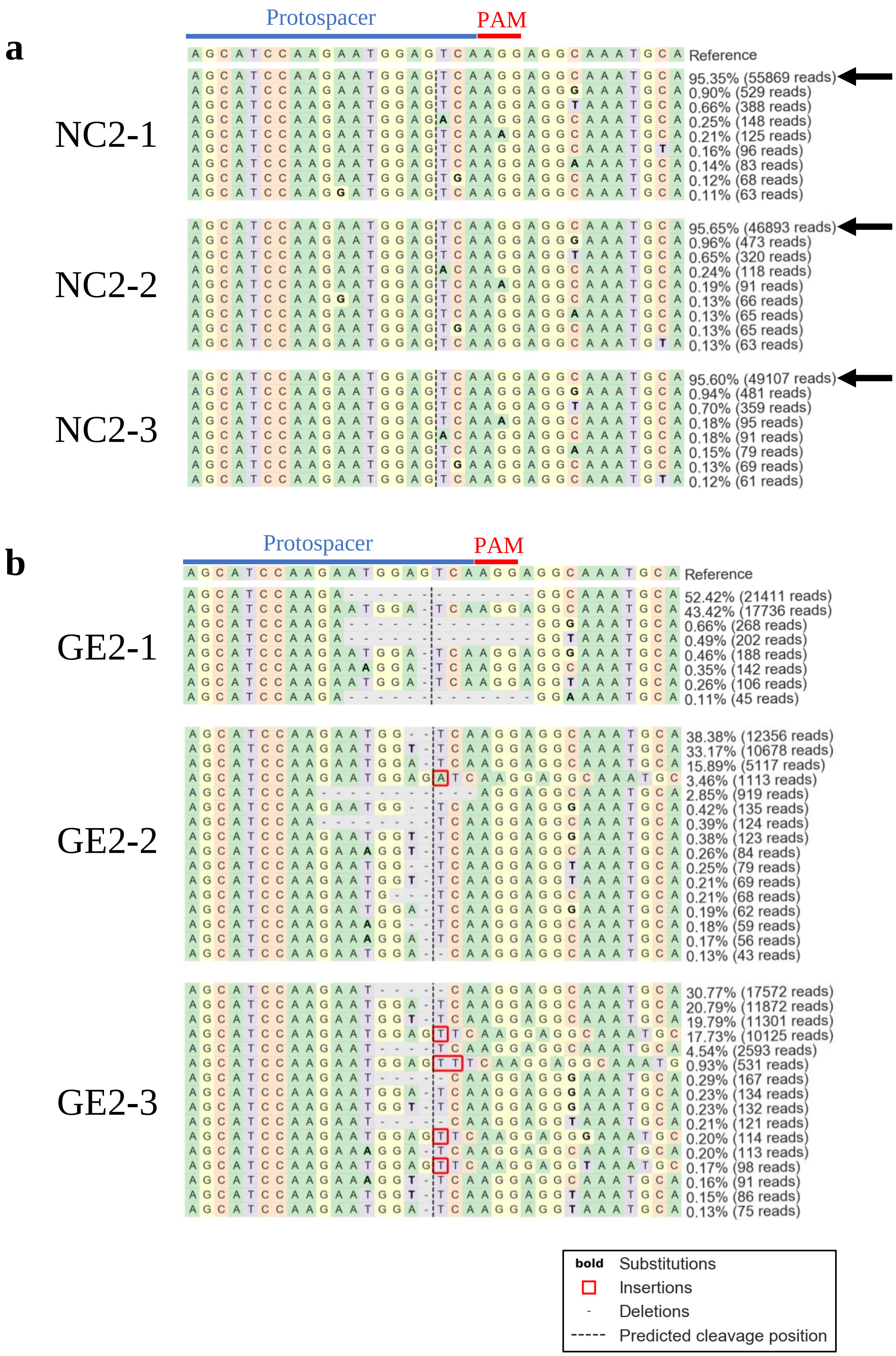


**Supplementary Figure S4**. Amplicon-sequencing analysis for on-target site in negative control (**a**) and genome-edited (**b**) rice callus cultivated for two months (n=3). The site that accumulated > 0.1% mutations are shown. The reads counts and its rate were displayed on each sequences.

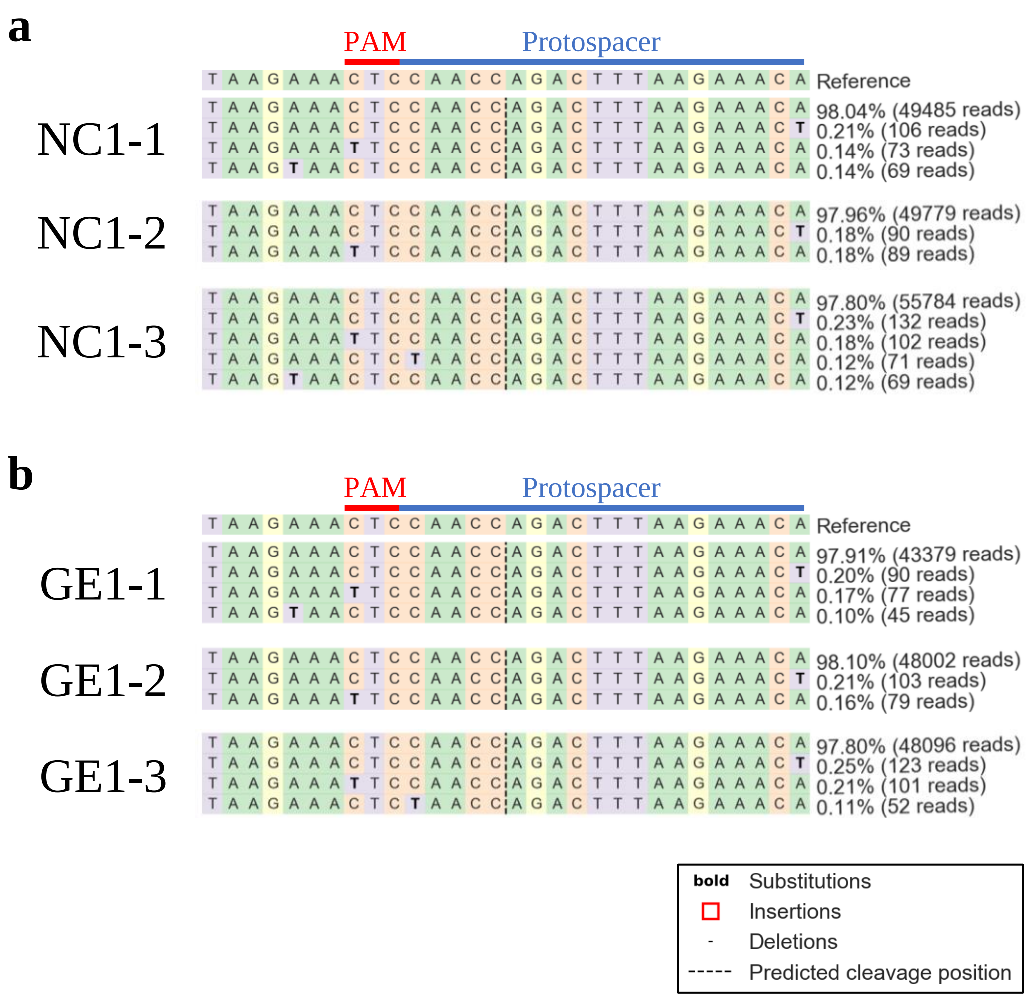


**Supplementary Figure S5**. Amplicon-sequencing analysis for off-target site 1 in negative control (**a**) and genome-edited (**b**) rice callus cultivated for one month (n=3). The site that accumulated > 0.1% mutations are shown. The reads counts and its rate were displayed on each sequences. The highly homologous sequence with the protospacer at the on-target site was shown as the predicted protospacer.

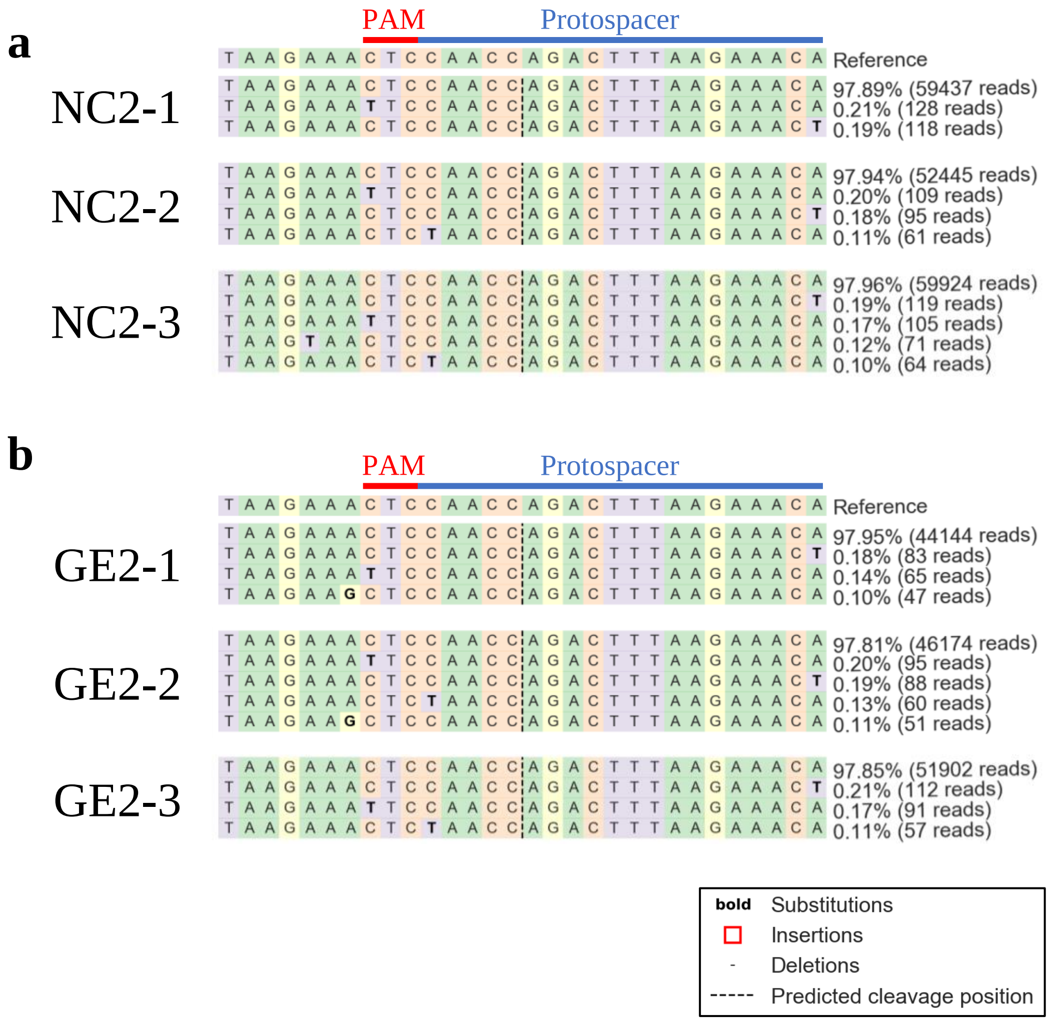


**Supplementary Figure S6**. Amplicon-sequencing analysis for off-target site 1 in negative control (**a**) and genome-edited (**b**) rice callus cultivated for two months (n=3). The site that accumulated > 0.1% mutations are shown. The reads counts and its rate were displayed on each sequences. The highly homologous sequence with the protospacer at the on-target site was shown as the predicted protospacer.

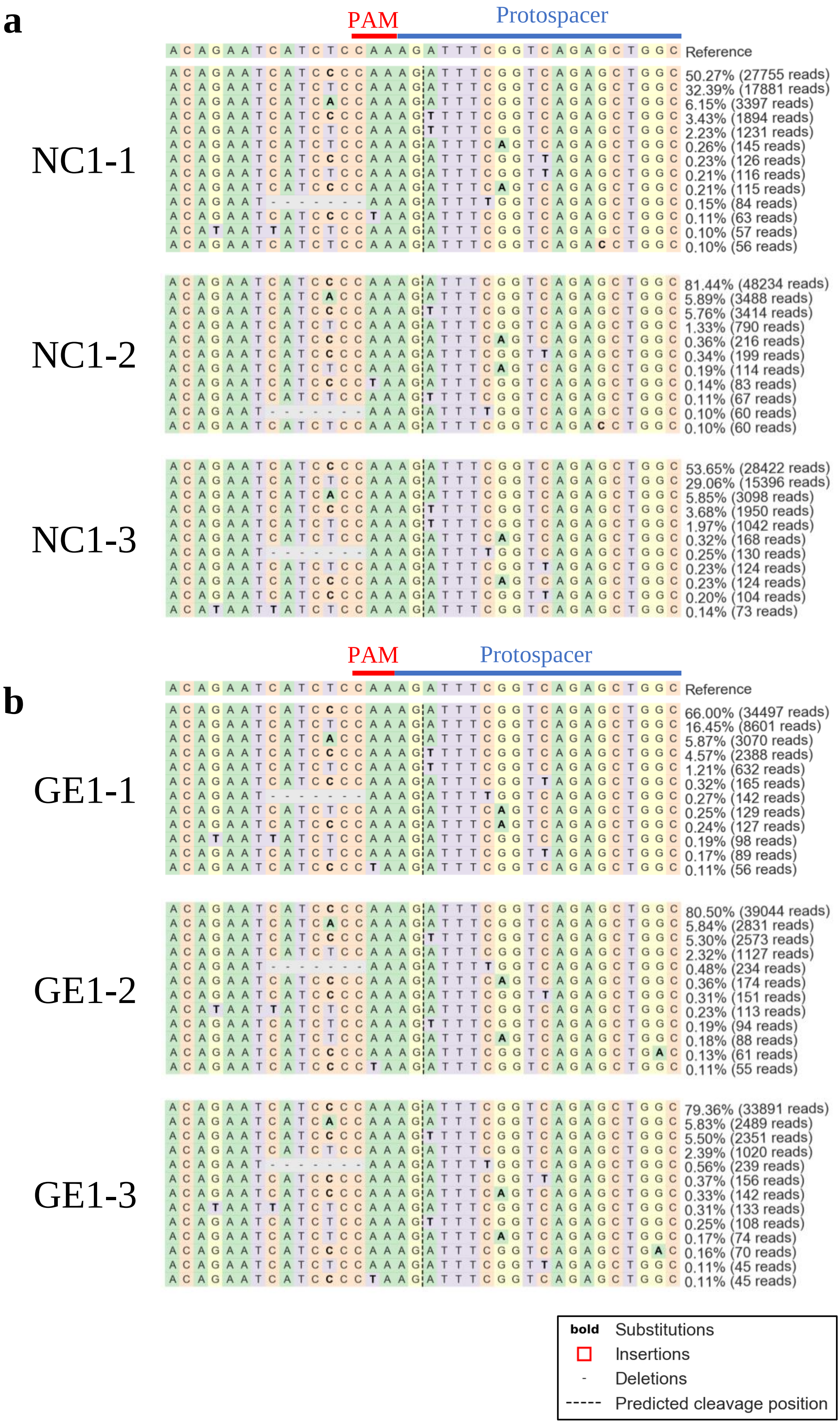


**Supplementary Figure S7**. Amplicon-sequencing analysis for off-target site 2 in negative control (**a**) and genome-edited (**b**) rice callus cultivated for one month (n=3). The site that accumulated > 0.1% mutations are shown. The reads counts and its rate were displayed on each sequences. The highly homologous sequence with the protospacer at the on-target site was shown as the predicted protospacer.

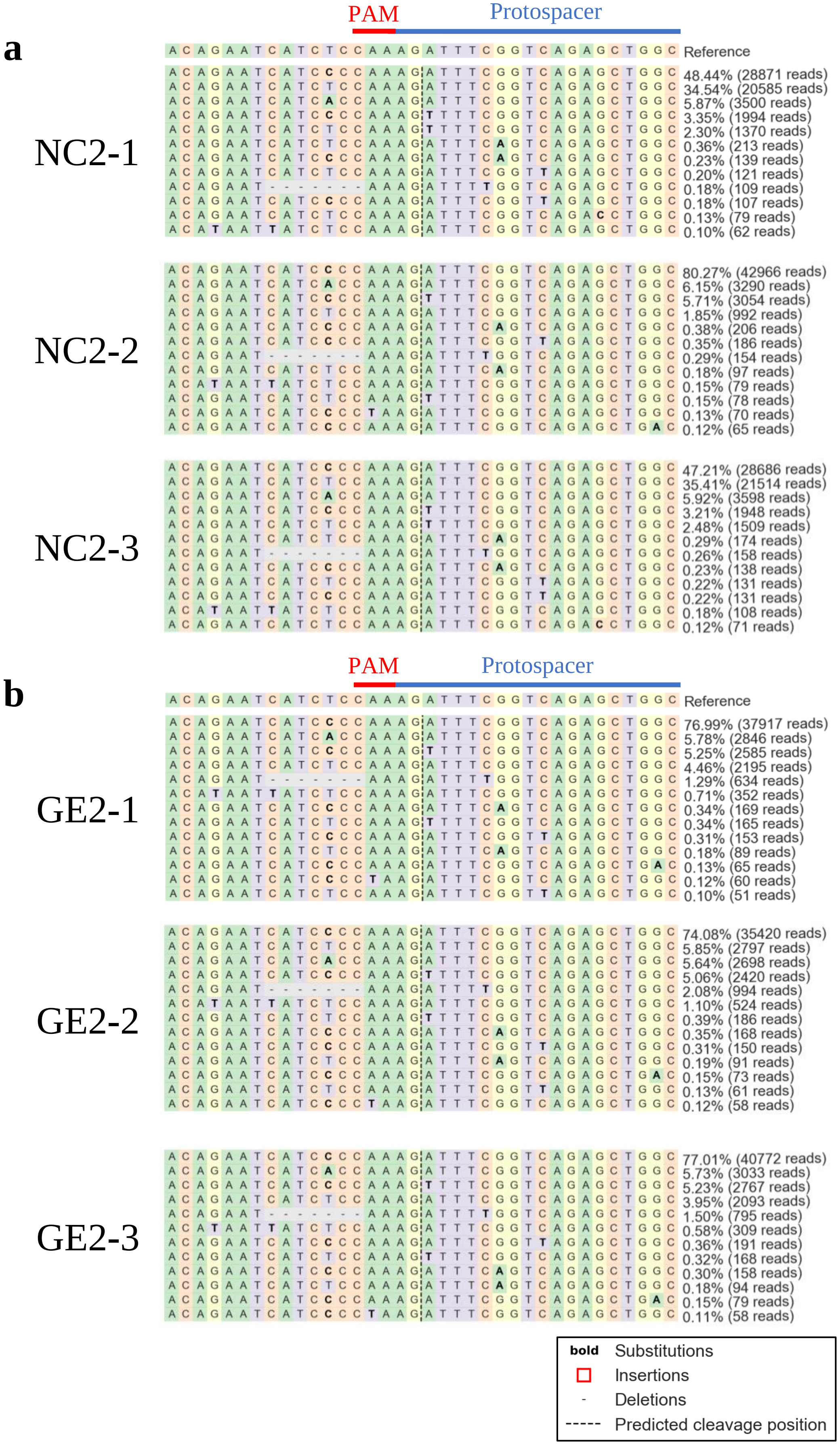


**Supplementary Figure S8**. Amplicon-sequencing analysis for off-target site 2 in negative control (**a**) and genome-edited (**b**) rice callus cultivated for two months (n=3). The site that accumulated > 0.1% mutations are shown. The reads counts and its rate were displayed on each sequences. The highly homologous sequence with the protospacer at the on-target site was shown as the predicted protospacer.

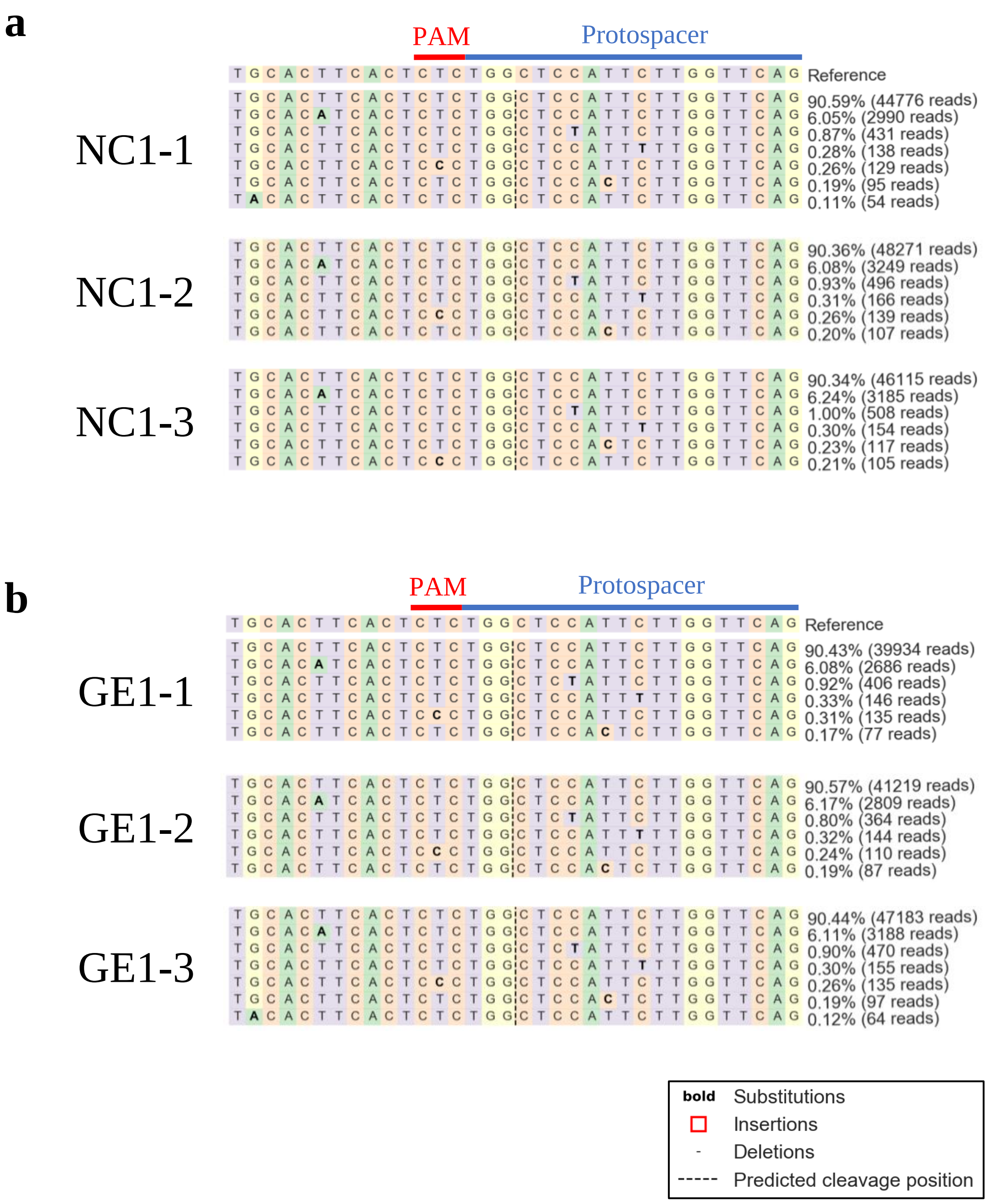


**Supplementary Figure S9**. Amplicon-sequencing analysis for off-target site 3 in negative control (**a**) and genome-edited (**b**) rice callus cultivated for one month (n=3). The site that accumulated > 0.1% mutations are shown. The reads counts and its rate were displayed on each sequences. The highly homologous sequence with the protospacer at the on-target site was shown as the predicted protospacer.

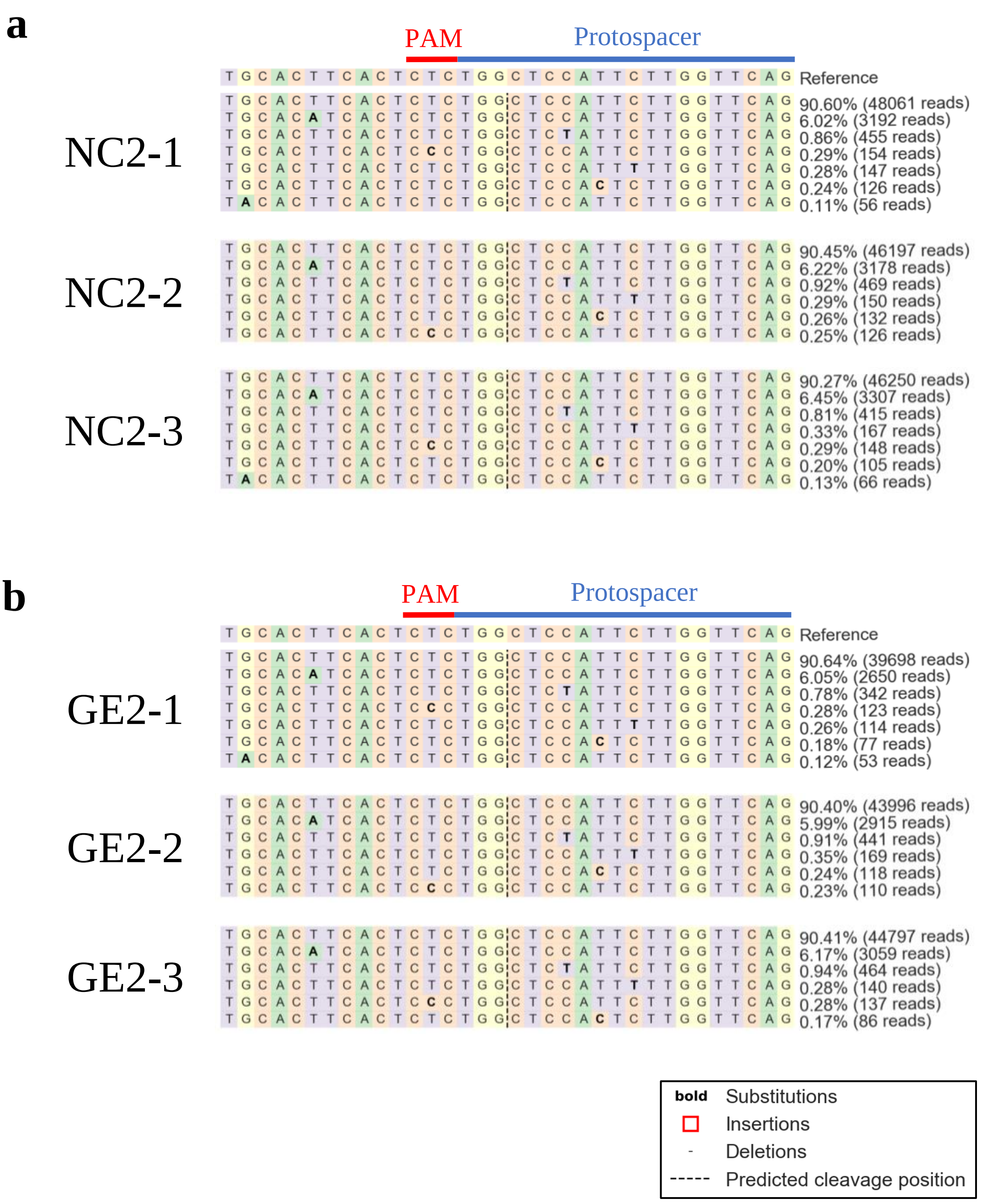


**Supplementary Figure S10**. Amplicon-sequencing analysis for off-target site 3 in negative control (**a**) and genome-edited (**b**) rice callus cultivated for two months (n=3). The site that accumulated > 0.1% mutations are shown. The reads counts and its rate were displayed on each sequences. The highly homologous sequence with the protospacer at the on-target site was shown as the predicted protospacer.

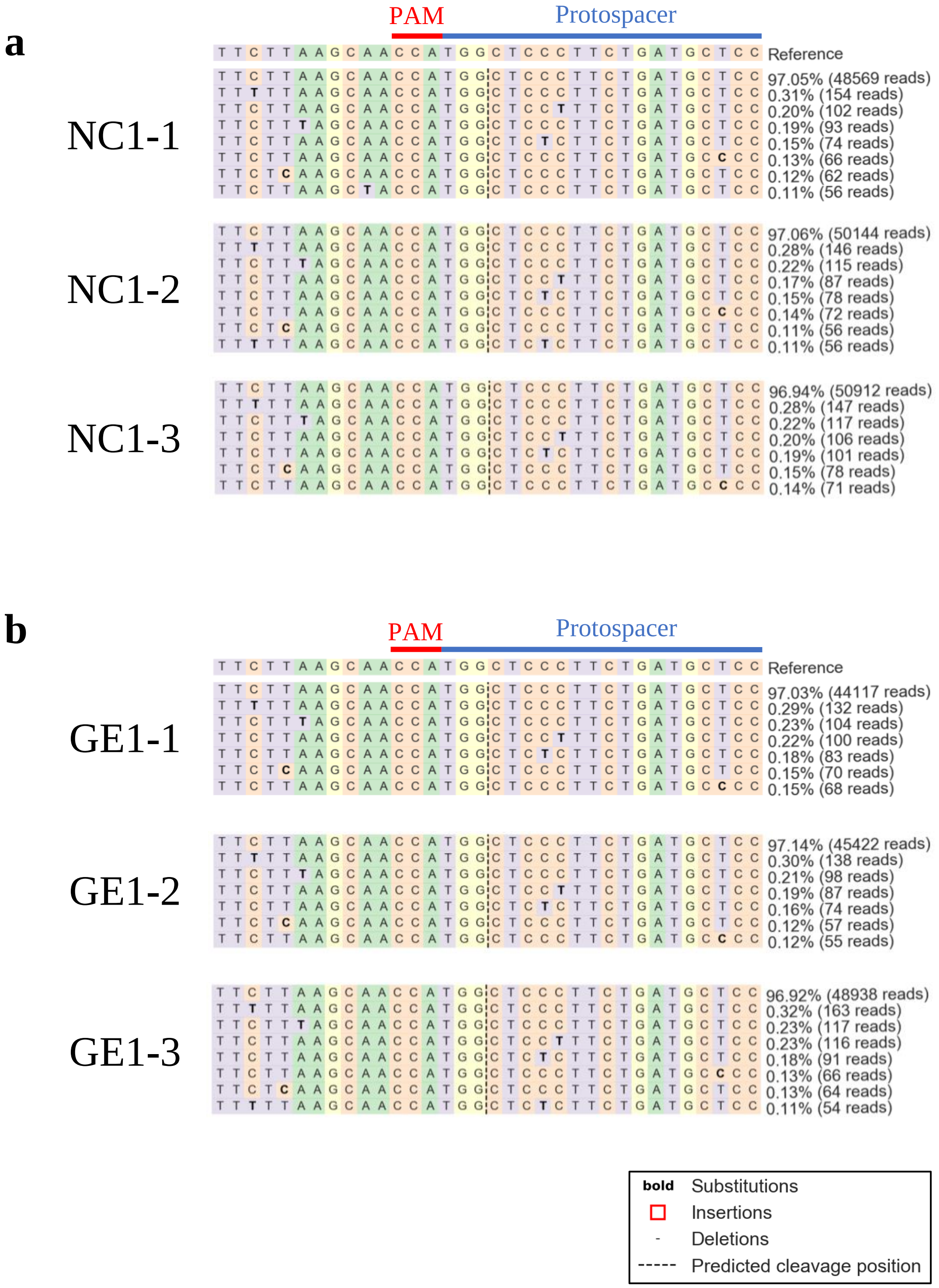


**Supplementary Figure S11.** Amplicon-sequencing analysis for off-target site 4 in negative control (**a**) and genome-edited (**b**) rice callus cultivated for one month (n=3). The site that accumulated > 0.1% mutations are shown. The reads counts and its rate were displayed on each sequences. The highly homologous sequence with the protospacer at the on-target site was shown as the predicted protospacer.

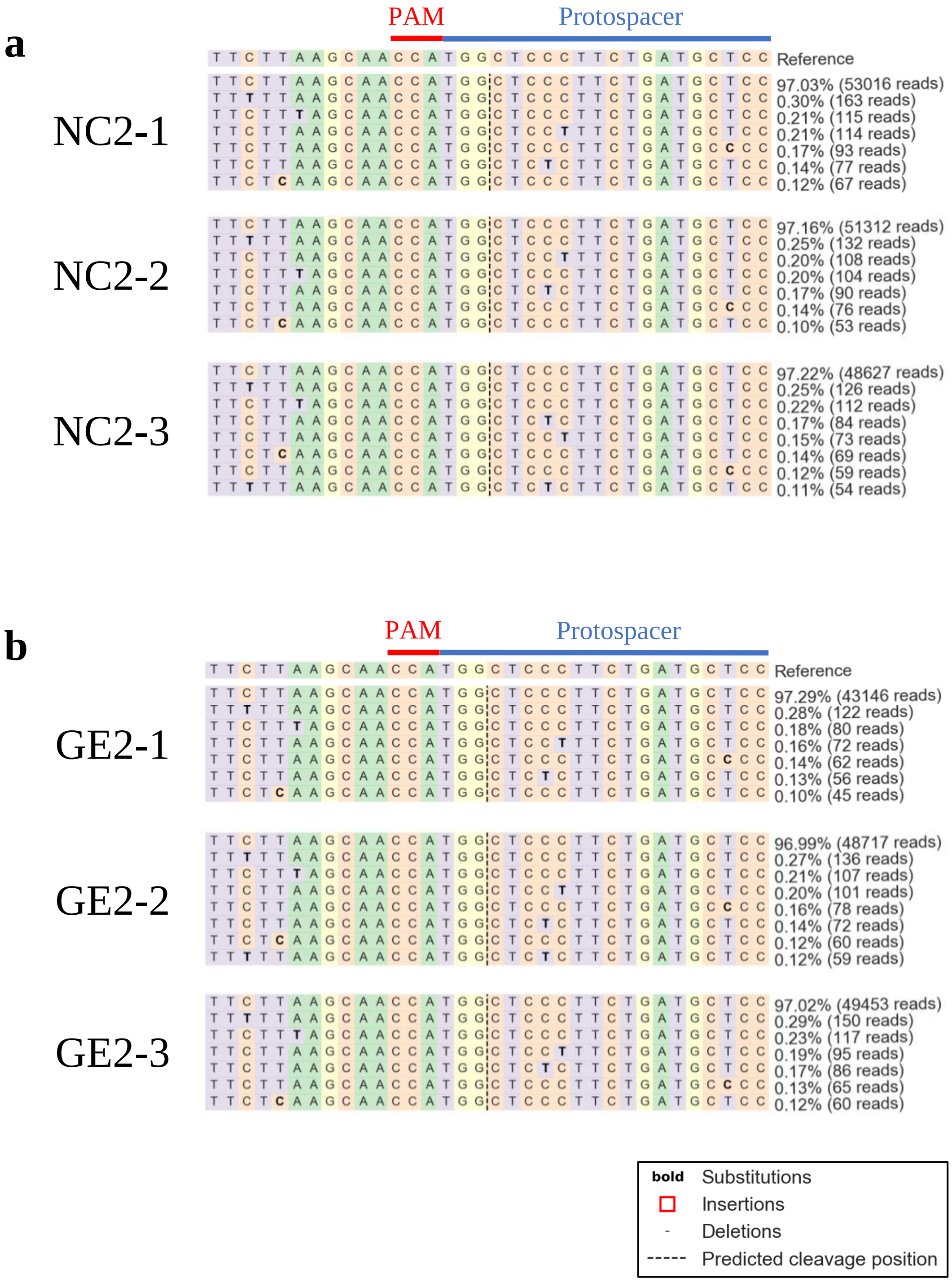


**Supplementary Figure S12.** Amplicon-sequencing analysis for off-target site 4 in negative control (**a**) and genome-edited (**b**) rice callus cultivated for two months (n=3). The site that accumulated > 0.1% mutations are shown. The reads counts and its rate were displayed on each sequences. The highly homologous sequence with the protospacer at the on-target site was shown as the predicted protospacer.

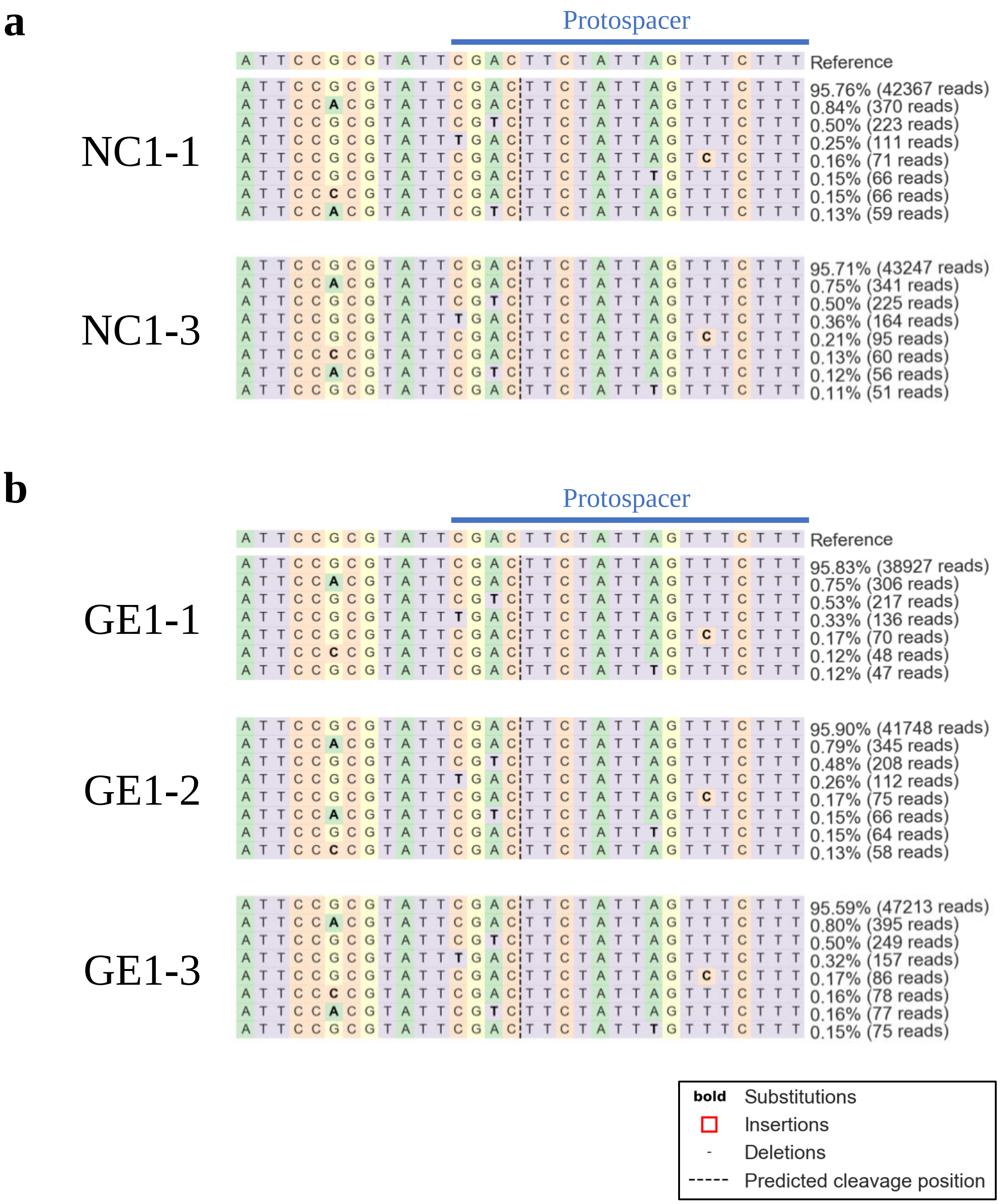


**Supplementary Figure S13.** Amplicon-sequencing analysis for off-target site 5 in negative control (n=2) (**a**) and genome-edited (n=3) (**b**) rice callus cultivated for one month. The site that accumulated > 0.1% mutations are shown. The reads counts and its rate were displayed on each sequences. The highly homologous sequence with the protospacer at the on-target site was shown as the predicted protospacer. There were no typical PAM around predicted protospacer sequence.

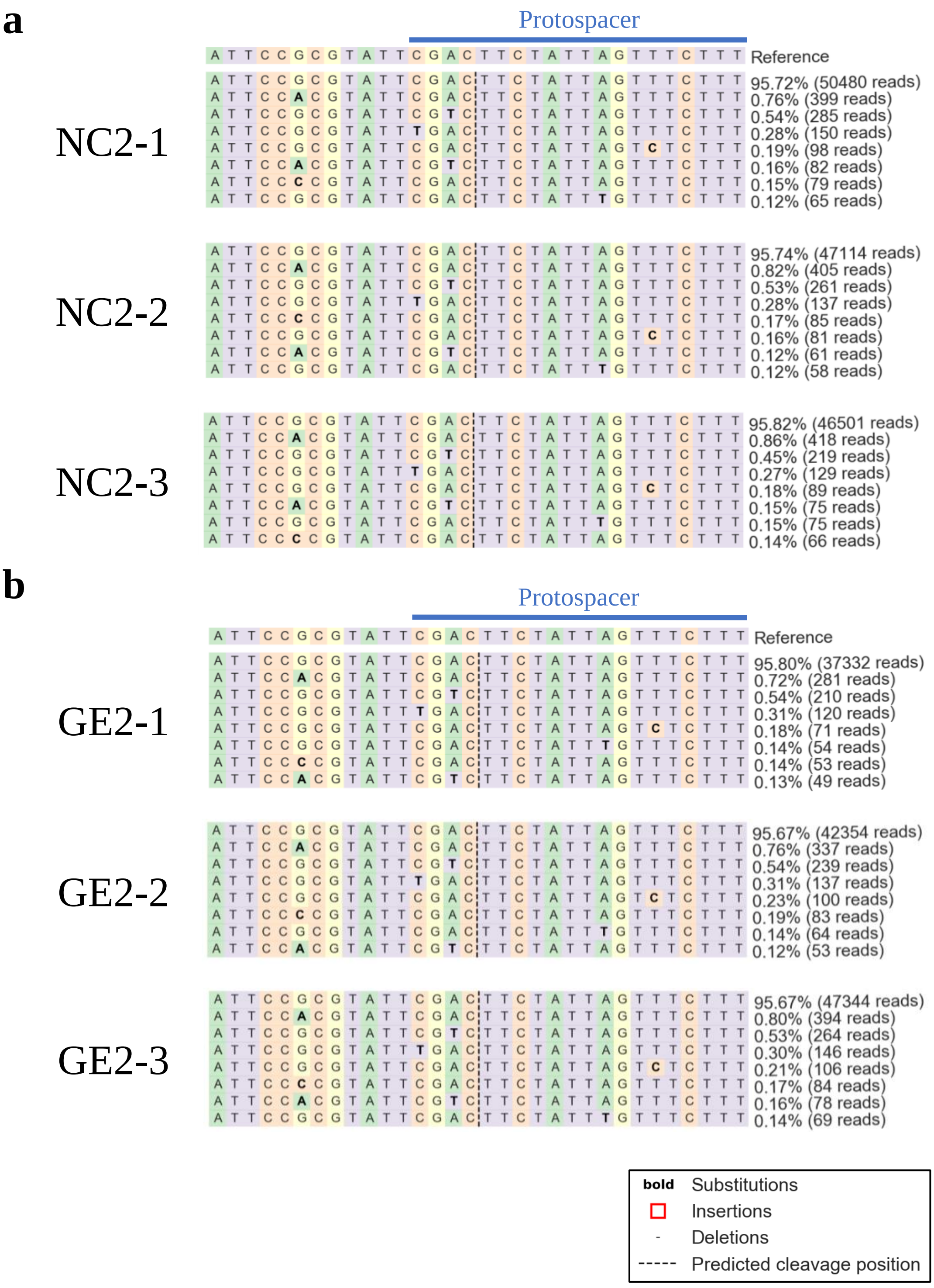


**Supplementary Figure S14.** Amplicon-sequencing analysis for off-target site 5 in negative control (**a**) and genome-edited (**b**) rice callus cultivated for two months (n=3). The site that accumulated > 0.1% mutations are shown. The reads counts and its rate were displayed on each sequences. The highly homologous sequence with the protospacer at the on-target site was shown as the predicted protospacer. There were no typical PAM around predicted protospacer sequences.

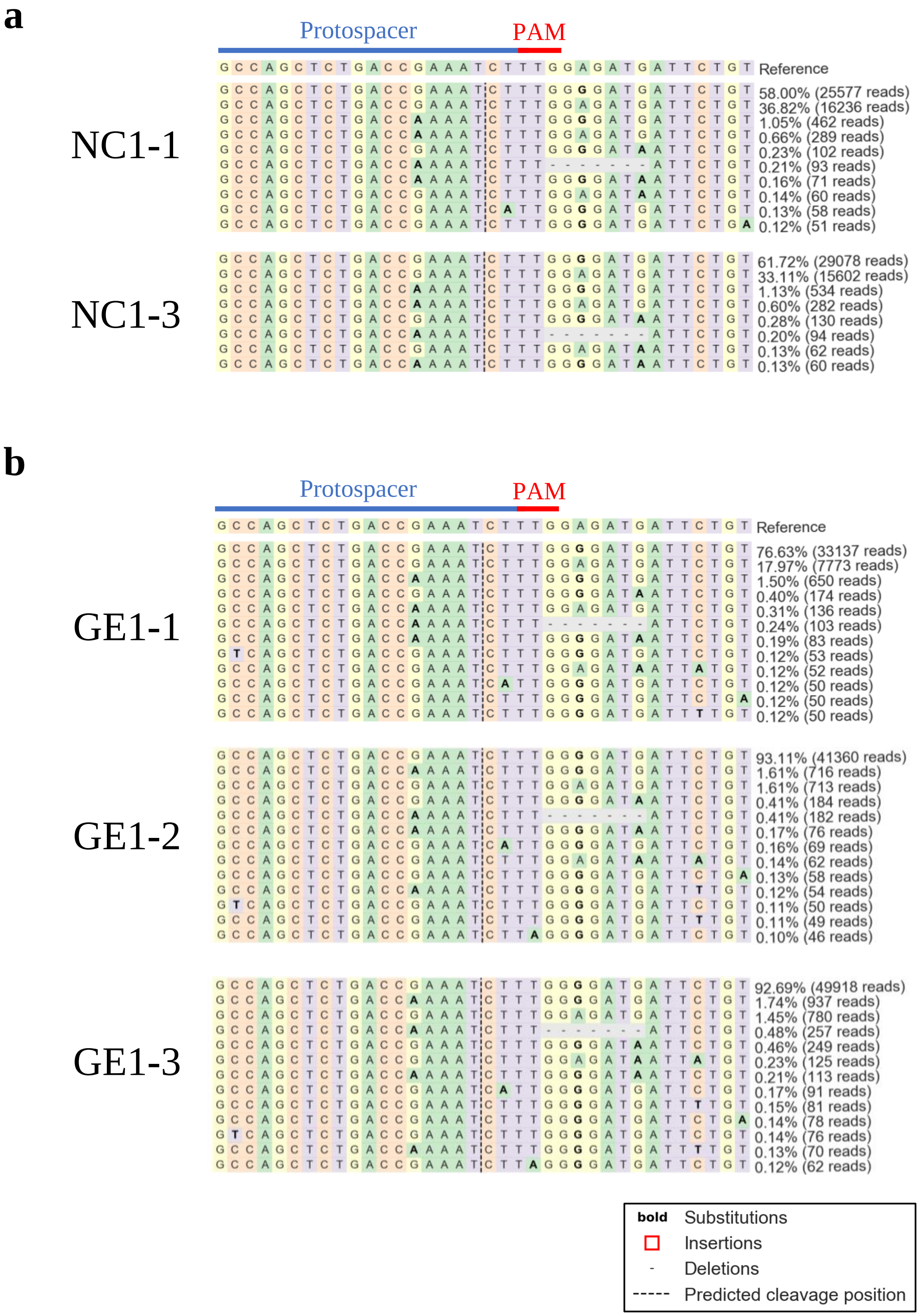


**Supplementary Figure S15.** Amplicon-sequencing analysis for off-target site 6 in negative control (n=2) (**a**) and genome-edited (n=3) (**b**) rice callus cultivated for one month. The site that accumulated > 0.1% mutations are shown. The reads counts and its rate were displayed on each sequences. The highly homologous sequence with the protospacer at the on-target site was shown as the predicted protospacer.

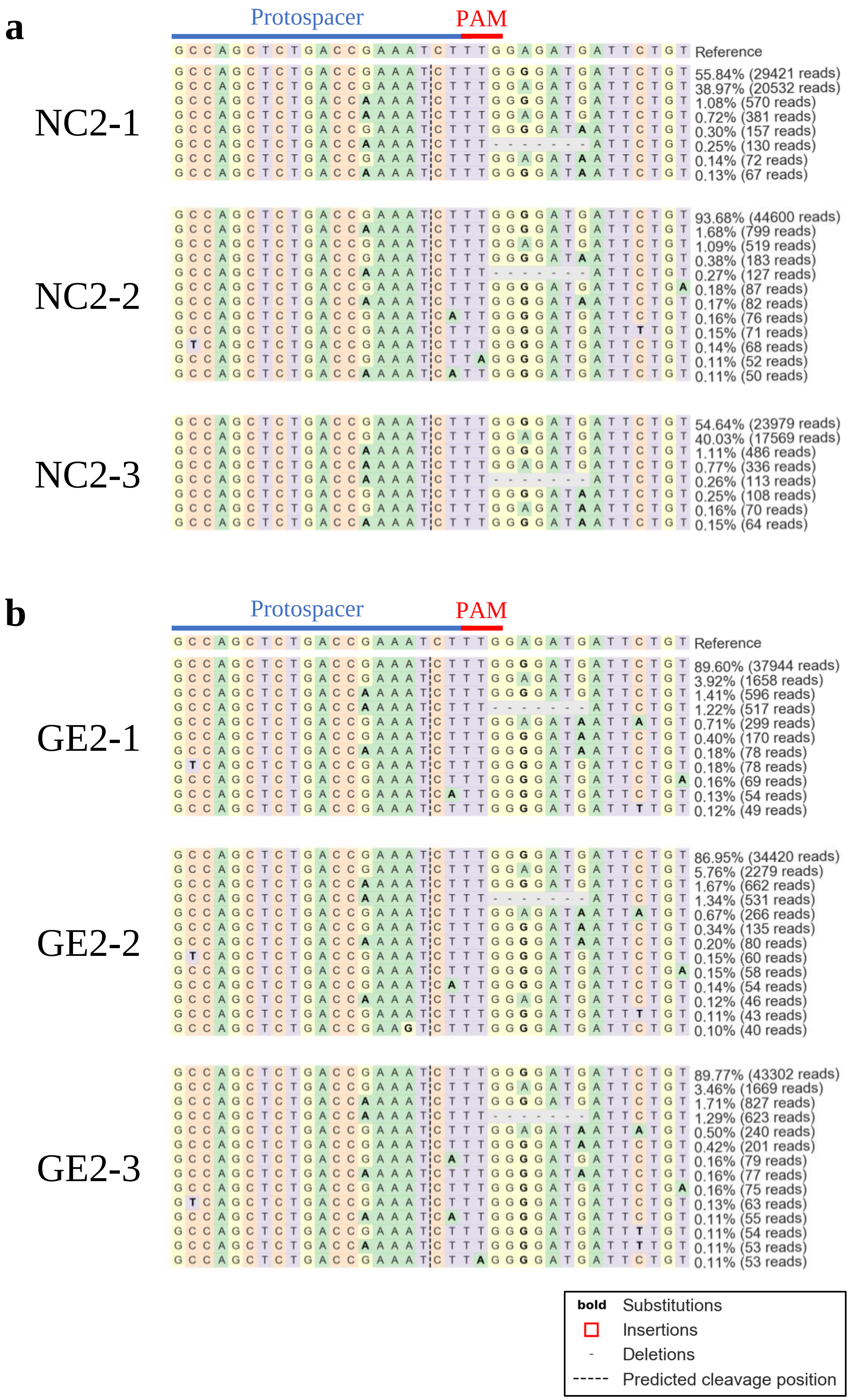


**Supplementary Figure S16.** Amplicon-sequencing analysis for off-target site 6 in negative control (**a**) and genome-edited (**b**) rice callus cultivated for two months (n=3). The site that accumulated > 0.1% mutations are shown. The reads counts and its rate were displayed on each sequences. The highly homologous sequence with the protospacer at the on-target site was shown as the predicted protospacer.

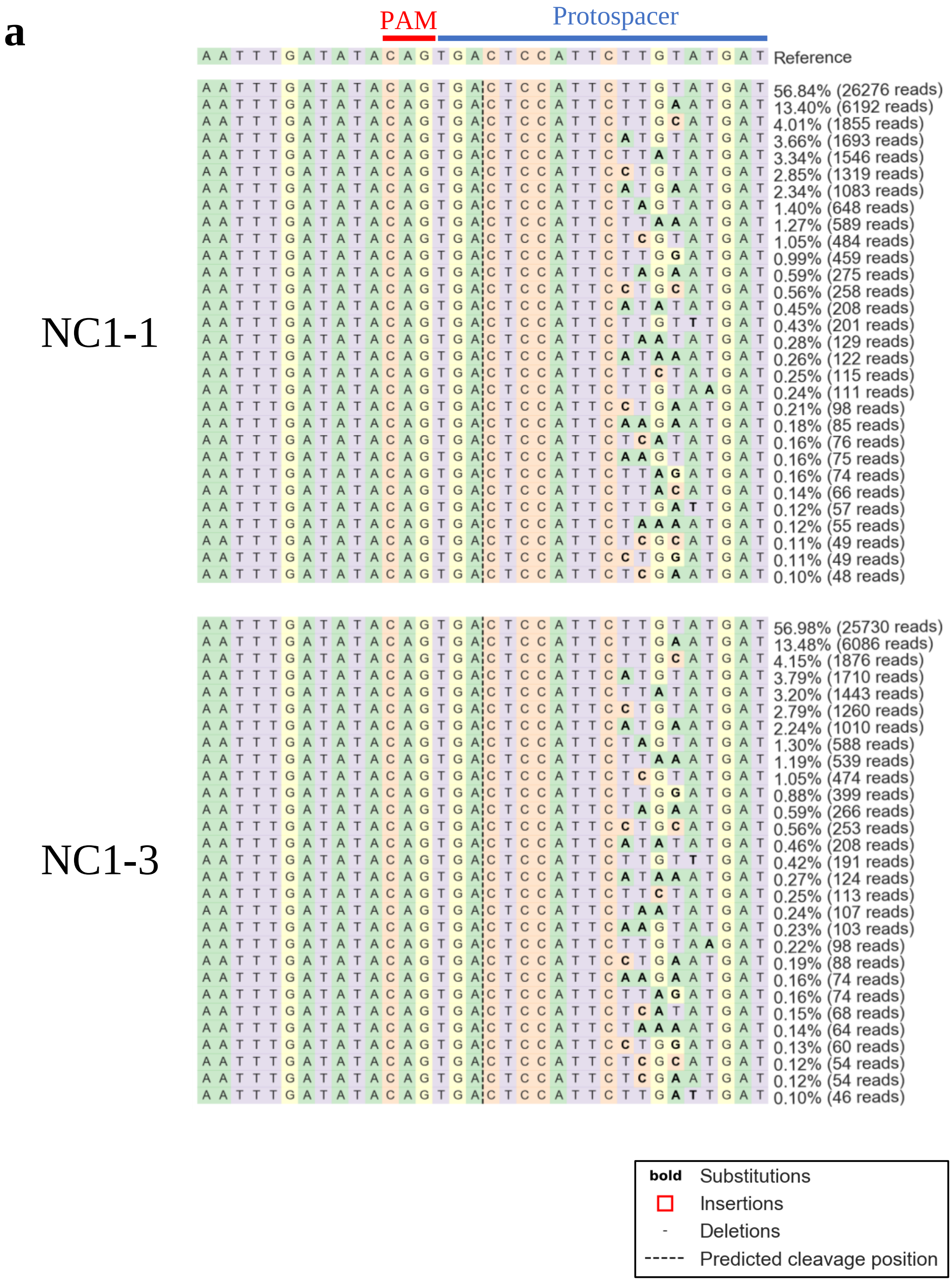


**Supplementary Figure S17**. Amplicon-sequencing analysis for off-target site 7 in negative control (n=2) (**a**) and genome-edited (n=3) (**b**) rice callus cultivated for one month. The site that accumulated > 0.1% mutations are shown. The reads counts and its rate were displayed on each sequences. The highly homologous sequence with the protospacer at the on-target site was shown as the predicted protospacer.

### Supplementary Figure S17. (continued)

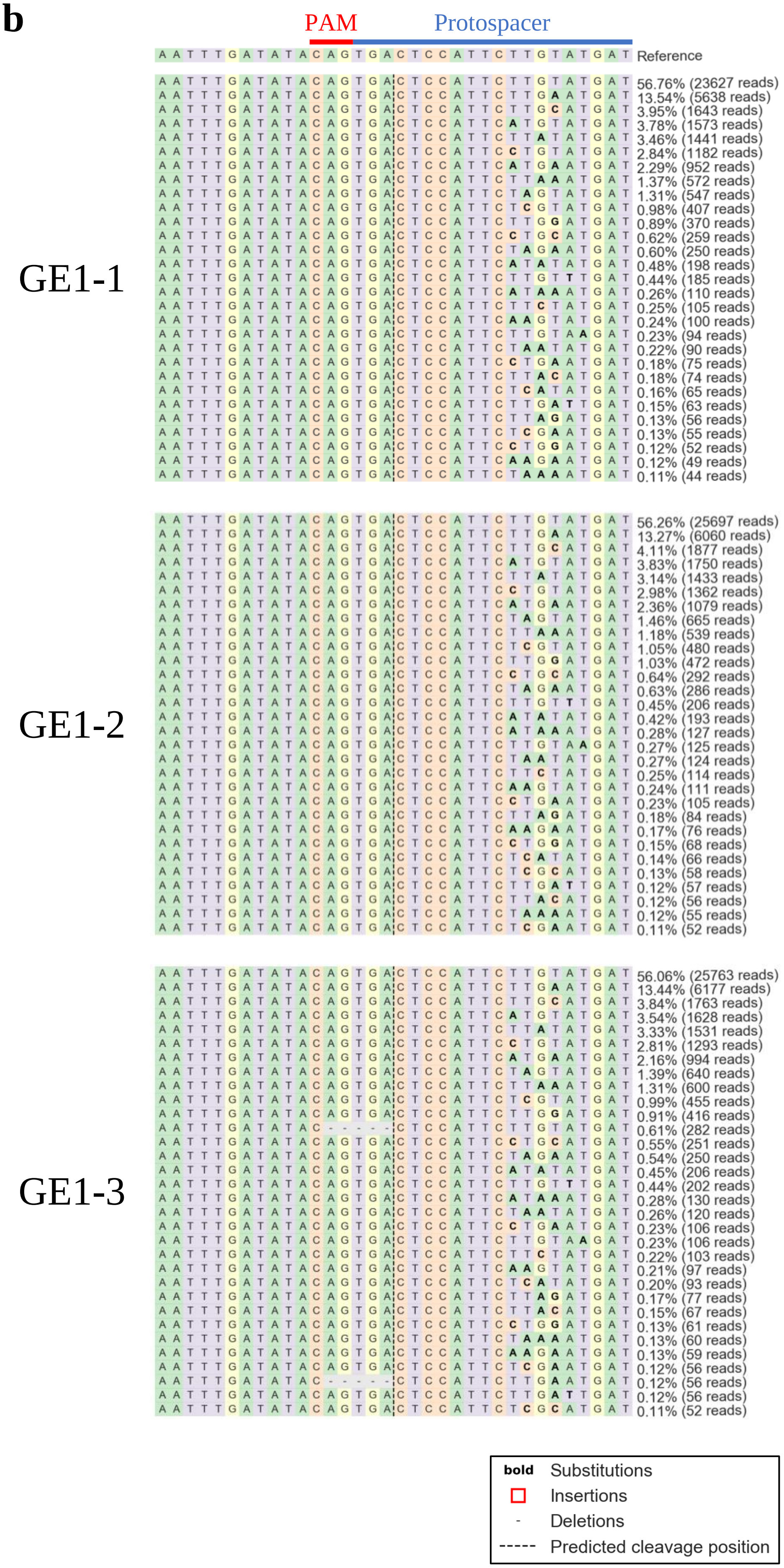

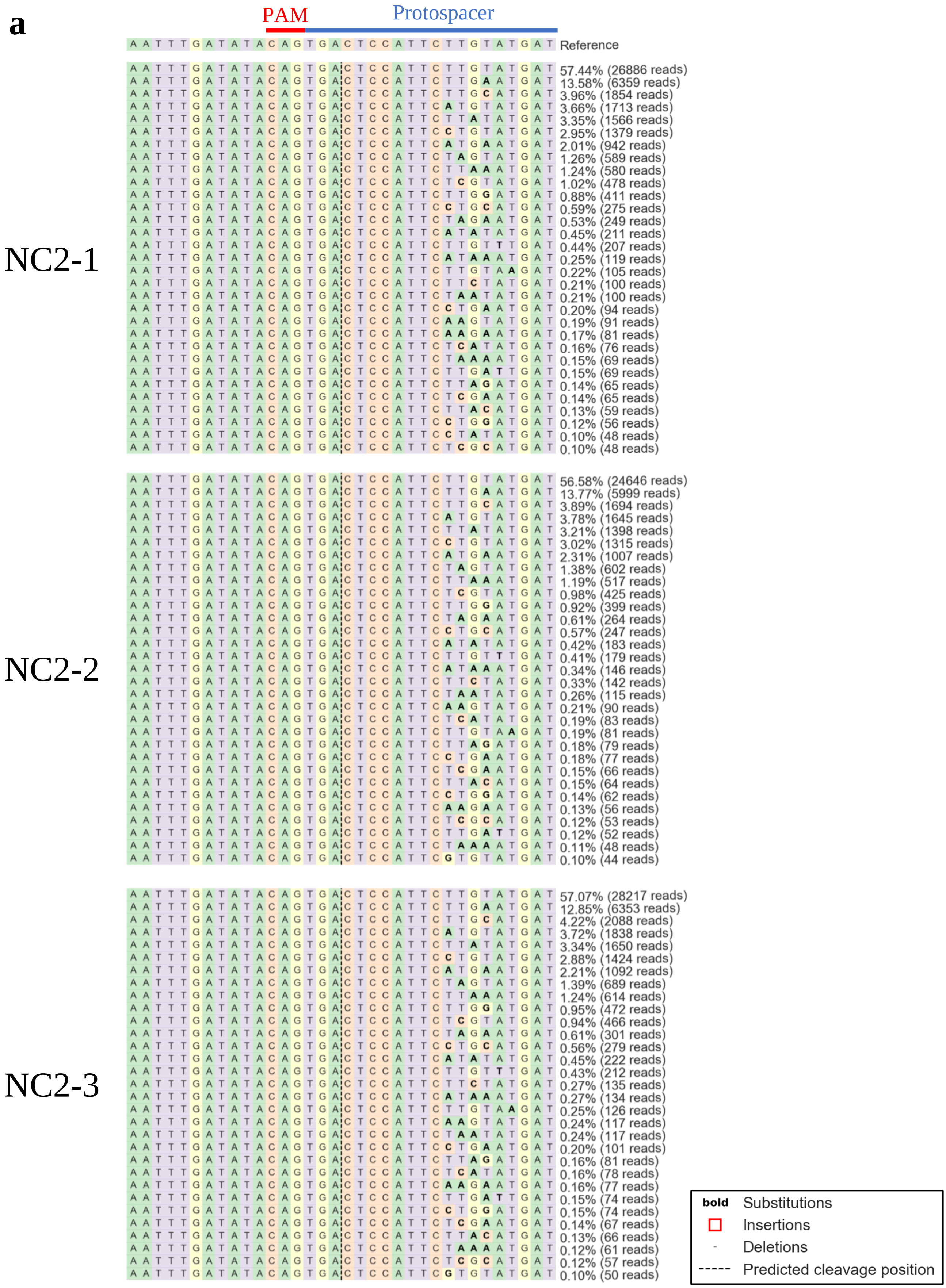


**Supplementary Figure S18**. Amplicon-sequencing analysis for off-target site 7 in negative control (**a**) and genome-edited (**b**) rice callus cultivated for two months (n=3). The site that accumulated > 0.1% mutations are shown. The reads counts and its rate were displayed on each sequences. The highly homologous sequence with the protospacer at the on-target site was shown as the predicted protospacer.

**Supplementary Figure S18. (continued)**

**b**

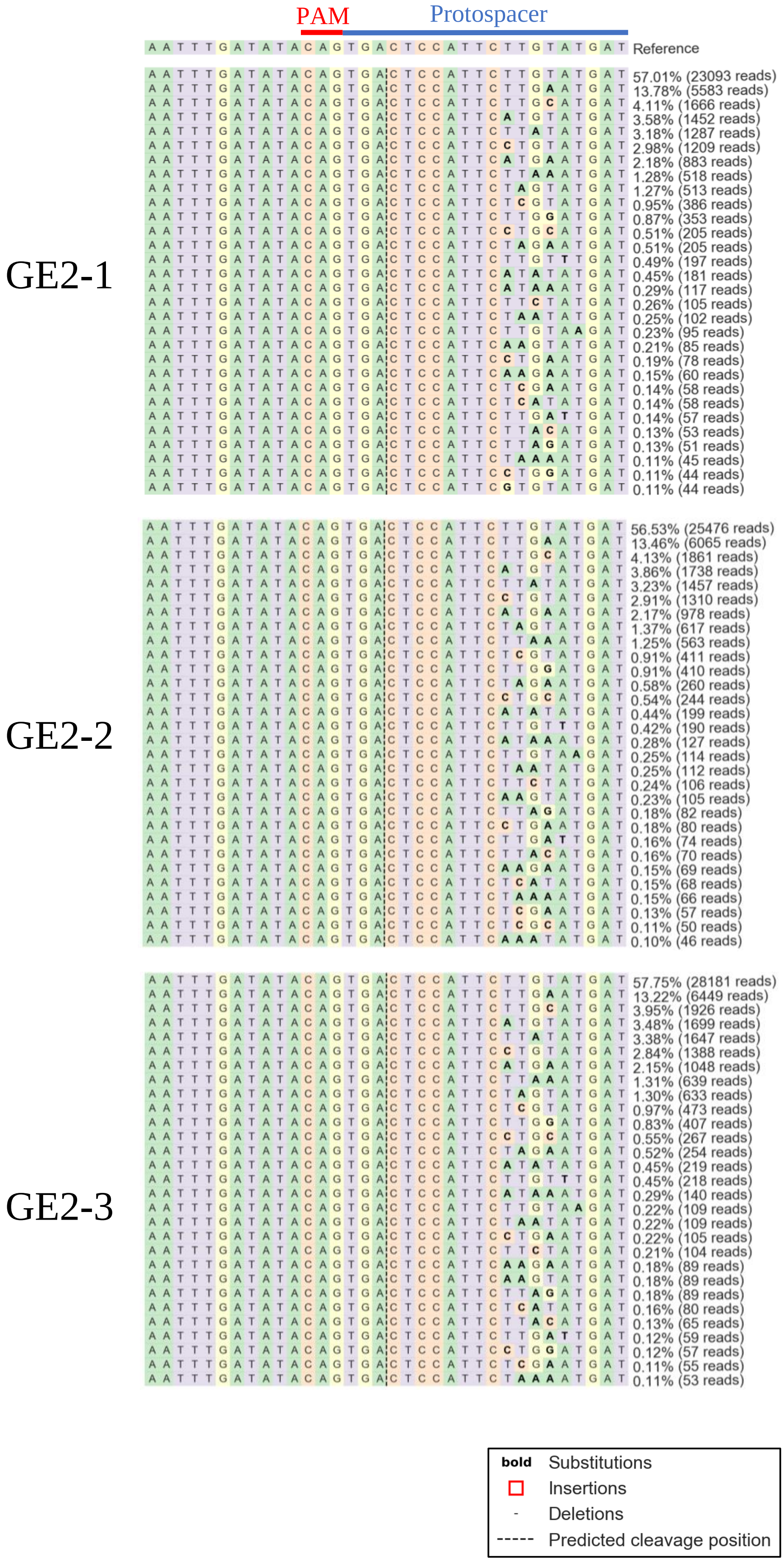

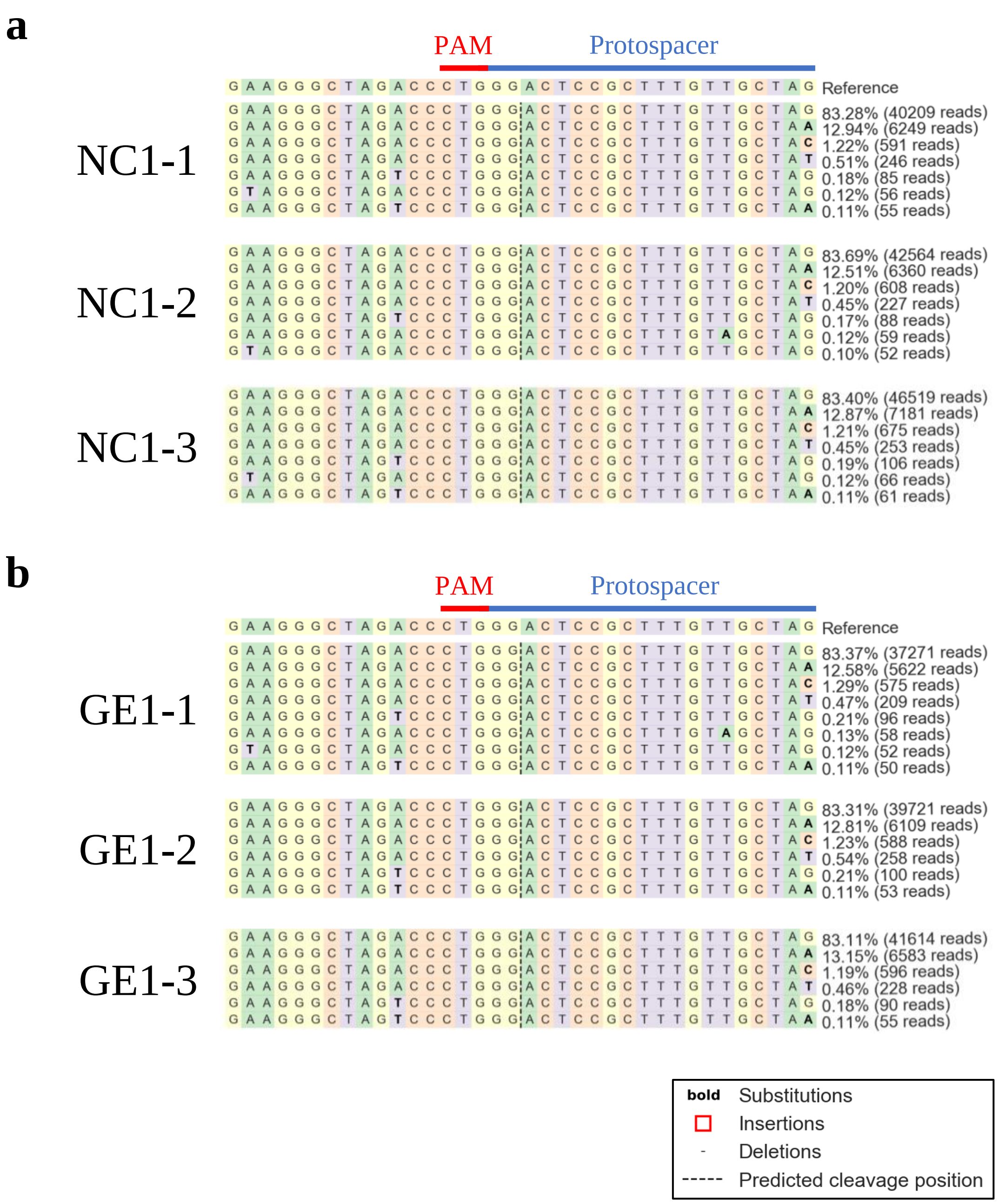


**Supplementary Figure S19**. Amplicon-sequencing analysis for off-target site 8 in negative control (**a**) and genome-edited (**b**) rice callus cultivated for one month (n=3). The site that accumulated > 0.1% mutations are shown. The reads counts and its rate were displayed on each sequences. The highly homologous sequence with the protospacer at the on-target site was shown as the predicted protospacer.

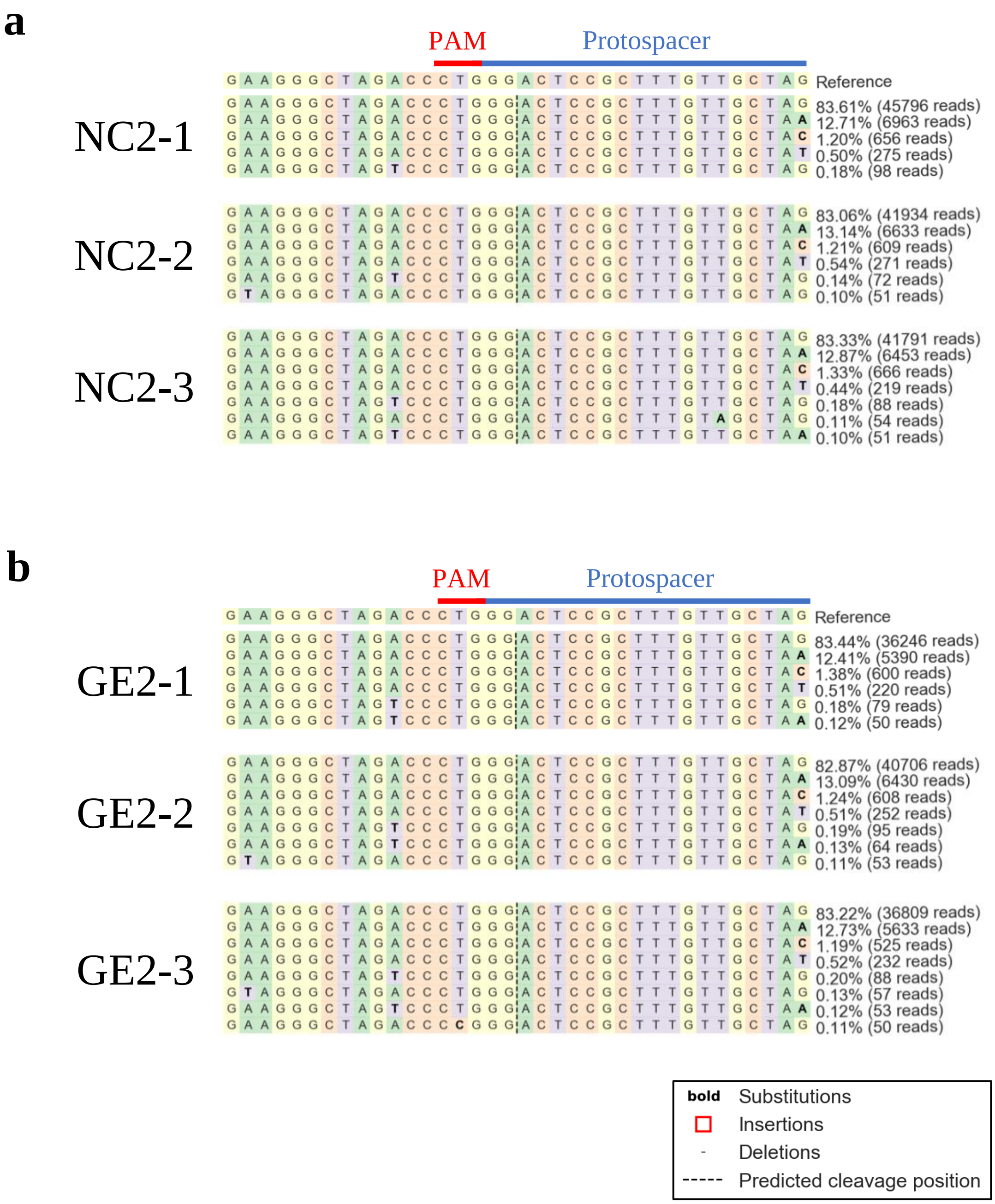


**Supplementary Figure S20**. Amplicon-sequencing analysis for off-target site 8 in negative control (**a**) and genome-edited (**b**) rice callus cultivated for two months (n=3). The site that accumulated > 0.1% mutations are shown. The reads counts and its rate were displayed on each sequences. The highly homologous sequence with the protospacer at the on-target site was shown as the predicted protospacer.
