## Supplementary Methods for "Strategy for detecting off-target sites in genome-edited rice"

Index

**Equipment**

**Reagents**

**SITE-Seq Procedure**

A. Purification of gDNA for plant (callus or seedling plant)

B. Amplification and purification of dsDNA template for gRNA synthesis

C. gRNA synthesis

D. Purification of the gRNA

E. Digestion of gDNA with RNP

F. Purification of RNP digested gDNA

G. Adapter 1 ligation

H. Fragmentation

I. Adapter 2 ligation

J. Affinity purification by streptavidin beads

K. Recovery PCR

L. Indexing PCR

M. Cleanup of the DNA library

N. Quantification and qualification of the DNA library for sequencing

O. Sequencing with Illumina MiSeq

**Amplicon-sequencing Procedure**

A. Amplification of the target sequences

B. Indexing PCR

C. DNA library QC

D. Sequencing with Illumina iSeq 100

**Equipment**

1. Mortar and pestle
2. Vortex Mixer Vortex-Genie 2 (Scientific Industries, Inc., Bohemia, NY, USA)
3. Water Bath Shaker PERSONAL-11 SD (TAITEC, Co., Saitama, Japan)
4. HulaMixer Sample Mixer (Invitrogen Co., Carlsbad, CA, USA)
5. Tube Centrifuge MX-305 (TOMY SEIKO CO., LTD., Tokyo, Japan)
6. Dry Thermo Unit DTU-1B (TAITEC, Co., Saitama, Japan)
7. Magnetic stand DynaMag- Spin Magnet (Invitrogen Co., Carlsbad, CA, USA)
8. Electrophoresis system Mupid-2plus (Mupid, Tokyo, Japan)
9. Spectrophotometer NanoDrop 1000 (Themo Fischer Scientific, Inc., Waltham, MA, USA)
10. Thermal Cycler Veriti 200 (Applied Biosystems, Waltham, MA, USA)
11. Precast agarose electrophoresis system E-Gel (Invitrogen Co., Carlsbad, CA, USA)
12. Qubit Fluorometer (Invitrogen Co., Carlsbad, CA, USA)
13. High-Resolution Automated Electrophoresis Agilent 2100 Bioanalyzer (Agilent Technologies, Inc., Santa Clara, CA USA)
14. NGS MiSeq System (Illumina, Inc., San Diego, CA, USA)
15. NGS iSeq 100 System (Illumina, Inc., San Diego, CA, USA)

**Reagents**

1. Hexadecyltrimethylammonium Bromide (FUJIFILM Wako Pure Chemical Co., Osaka, Japan)
2. 1 mol/l Tris-HCl Buffer Solution (pH 8.0) (NACALAI TESQUE, INC., Kyoto, Japan)
3. UltraPure™ 0.5 M EDTA, pH 8.0 (Invitrogen Co., Carlsbad, CA, USA)
4. Sodium Chloride (NACALAI TESQUE, INC., Kyoto, Japan)
5. Distilled Water (NIPPON GENE CO., LTD., Tokyo, Japan)
6. Chloroform (FUJIFILM Wako Pure Chemical Co., Osaka, Japan)
7. 3-Methyl-1-butanol (Sigma-Aldrich, Co., LLC., St Lois, MO, USA)
8. 2-Propanol (FUJIFILM Wako Pure Chemical Co., Osaka, Japan)
9. Ethanol (FUJIFILM Wako Pure Chemical Co., Osaka, Japan)
10. RNase A (100 mg/ml) (NIPPON GENE CO., LTD., Tokyo, Japan)
11. SPRIselect (Beckman Coulter, Inc., Brea, CA, USA)
12. RNA secure (Invitrogen Co., Carlsbad, CA, USA)
13. Herculase II Fusion DNA polymerase (Agilent Technologies, Inc., Santa Clara, CA USA)
14. NucleoSpin Gel and PCR Clean-up (MACHEREY-NAGEL, GmbH & Co. KG, Dueren,Germany)
15. SureGuide gRNA Synthesis Kit (Agilent Technologies, Inc., Santa Clara, CA USA)
16. GeneArt Platinum Cas9 Nuclease (Invitrogen Co., Carlsbad, CA, USA)
17. HEPES (DOJINDO LABORATORIES, Kumamoto, Japan)
18. Potassium Chloride (FUJIFILM Wako Pure Chemical Co., Osaka, Japan)
19. Magnesium Chloride Hexahydrate (FUJIFILM Wako Pure Chemical Co., Osaka, Japan)
20. Glycerol (Sigma-Aldrich, Co., LLC., St Lois, MO, USA)
21. Proteinase K (FUJIFILM Wako Pure Chemical Co., Osaka, Japan)
22. 3 M Sodium Acetate (pH 5.2) (NIPPON GENE CO., LTD., Tokyo, Japan)
23. dATP Solution (New England Biolabs Inc., Ipswich, MA, USA)
24. Klenow Fragment (3'-5' exo-) (New England Biolabs Inc., Ipswich, MA, USA)
25. T4 DNA Ligase (New England Biolabs Inc., Ipswich, MA, USA)
26. NEBNext dsDNA Fragmentase (New England Biolabs Inc., Ipswich, MA, USA)
27. NEBNext Ultra End Repair/ dA-tailing module（New England Biolabs Inc., Ipswich, MA, USA)
28. Blunt/TA Ligase Master Mix (New England Biolabs Inc., Ipswich, MA, USA)
29. Dynabeads M-280 streptavidin (Invitrogen Co., Carlsbad, CA, USA)
30. Phusion High-Fidelity PCR Master Mix with HF Buffer (New England Biolabs Inc., Ipswich, MA, USA)
31. Qubit dsDNA HS Assay Kit (Invitrogen Co., Carlsbad, CA, USA)

32. Agilent 2100 Bioanalyzer High Sensitivity DNA kit (Agilent Technologies, Inc., Santa Clara, CA USA)
33. Tween 20 (Sigma-Aldrich, Co., LLC., St Lois, MO, USA)
34. 1 mol/l Sodium Hydroxide Solution (NACALAI TESQUE, INC., Kyoto, Japan)
35. Lens cleaning tissue (Whatman plc, Kent, UK)
36. MiSeq Reagent Kit v3 (150 cycles) (Illumina, Inc., San Diego, CA, USA)
37. iSeq 100 i1 Reagent (300 Cycles) (Illumina, Inc., San Diego, CA, USA)

**SITE-Seq Procedure**

**A. Purification of gDNA for plant (callus or seedling plant)**

1. Prepare 1.5 × CTAB (Cetyl trimethyl ammonium bromide) buffer and CIA according to the following.

1.5 × CTAB buffer

| | |
|---|---|
| CTAB | 15 g |
| 1 M Tris-HCl, pH 8.0 | 75 mL |
| 0.5 M EDTA, pH 8.0 | 30 mL |
| NaCl | 61 g |
| DW | Fill up to 1 L |

CIA

| | |
|---|---|
| Chloroform | 96 mL |
| Isoamyl alcohol | 4 mL |

2. Grind up callus or seedling plant in liquid nitrogen.
3. Transfer the powdered sample into a 50 mL tube.
4. Add 2.5 mL of 1.5 × CTAB buffer per 1 g of sample.
5. Incubate at 56°C for 20 min with vortexing appropriately.
6. Add 1 × volumes of CIA (2.5 mL/g sample).
7. Mix using sample mixer at room temperature (RT) for 20 min.
8. Centrifuge at 7,000 × *g* and RT for 15 min.
9. Transfer the supernatant from aqueous layer to a new tube.
10. Add 0.7 × volumes of cold isopropanol and invert to mix.
11. Centrifuge at 10,000 × *g* and 4°C for 10 min.
12. Discard the supernatant, add appropriated cold 70% EtOH.
13. Centrifuge at 10,000 × *g* and 4°C for 10 min.
14. Discard EtOH entirely, resuspend the gDNA with 100 µL of DW.
15. Add 10 µL of DNase free-RNase A (10 mg/mL), and then incubate at 37°C for 30 min.
16. Add 0.04 × volumes of RNA secure (4.4 µL), incubate at 60°C for 10 min, and then cool to RT.
17. Add 0.4 × volumes of SPRIselect (46 µL), mix by pipetting, and then incubate at RT for 5 min.
18. Discard the supernatant using magnetic stand, and wash the beads with 180 µL of cold 85% EtOH three times.
19. Resuspend the beads in 50 µL of DW or TE, and incubate at RT for 5 min.
20. Collect the clear supernatant using magnetic stand, and perform agarose gel electrophoresis.

If necessary, repeat 0.4 × SPRIselect to remove sheared DNA or residual RNA (see Figure).

21. Measure the concentration of gDNA by NanoDrop 1000.

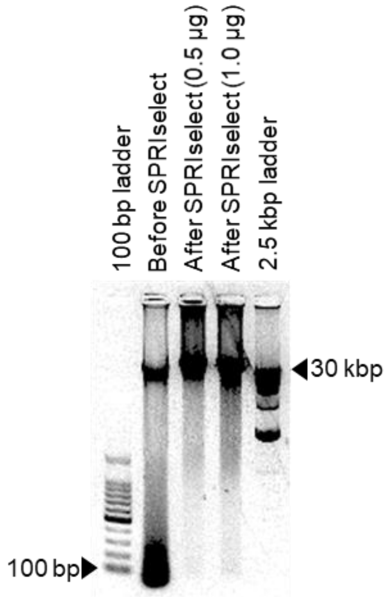


Figure. Agarose gel electrophoresis of isolated gDNA from rice callus by CTAB method with or without SPRIselect treatment.

**B. Amplification and purification of dsDNA template for gRNA synthesis**

* “gg” at the 5’ end of a guide is required for efficient transcription by T7 RNA polymerase.

1. Synthesize two oligos as follows (replace 20 nt of x to designed gRNA sequence, not include PAM. Fwd and Rev oligos were annealed between underlined nucleotides. y was indicated complemented nucleotide of x. e.g. x = A, y = T).

| Name | Sequence (5’-3’) |
|---|---|
| Oligo-Fwd | 5’-cgatgtaatacgactcactataggxxxxxxxxxxxxxxxxxxxxgttttagagctatgctgaaa-3’ |
| Oligo-Rev | 5’-aagcaccgactcggtgccactttttcaagttgataacggactagccttattttaacttgctatgctttcagcatagctctaaaacy-3’ |

2. Prepare the reaction mixture.

| | |
|---|---|
| 5 × Herculase II reaction buffer (Agilent) | 10 µL |
| dNTPs (2.5 mM each) | 4 µL |
| Oligo-Fwd (10 µM) | 5 µL |
| Oligo-Rev (10 µM) | 5 µL |
| Herculase II Fusion DNA polymerase | 1 µL |
| DW | 25 µL |
| Total | 50 µL |

3. Incubate at 95°C for 2 min → 60°C for 1 min → 72°C for 3 min.
4. Purify the DNA fragment from the agarose gel slice using NucleoSpin Gel and PCR Clean-up.
5. Measuring the concentration of dsDNA by NanoDrop 1000.

**C. gRNA synthesis**

1. Prepare the reaction mixture on ice (SureGuide gRNA synthesis kit).

| | |
|---|---|
| DEPC water | 7.5 μL |
| 5 × Transcription buffer | 5 μL |
| rATP | 1 μL |
| rCTP | 1 μL |
| rGTP | 1 μL |
| rUTP | 1 μL |
| 0.75 M DTT | 1 μL |
| Yeast Pyrophosphatase | 0.5 μL |
| RNase Block | 1 μL |
| T7 RNA polymerase | 1 μL |
| Total | 20 μL |

2. Add 5 μL of DNA template (1 μM) to the reaction mixture and incubate at 37°C for 16 h.
3. Add 1 μL of RNase-free DNase, mix well, and incubate for 15-20 min at 37°C.

**D. Purification of gRNA**

1. Seat a spin cup filter in a new 2 mL receptacle tube.
2. Add 200 μL of gRNA binding buffer to the reaction mixture and mix by pipetting.
3. Transfer the entire volume of sample (226 μL) to a spin cup filter.
4. Centrifuge the spin cup filter at maximum speed for 1 min.
5. Discard the elute, add 600 μL of gRNA wash buffer to the spin cup filter.
6. Centrifuge the spin cup filter at maximum speed for 1 min.
7. Discard the elute, centrifuge at maximum speed for 2 min to dry the filter matrix.
8. Transfer the filter cup to a new 1.5 mL tube.
9. Add 30 μL of gRNA elute buffer to the center of the filter.
10. Centrifuge at maximum speed for 1 min.
11. Measuring the concentration of gRNA by NanoDrop 1000.
12. Calculate the concentration (μM) = x (ng/μL) / [108 nt × 330 (g/mol)] × $10^3$
13. Store at -80°C until use.

**E. gDNA digestion with RNP**

* MW of SpCas9 is approximately 158,441 g/mol.

* MW of gRNA (108 nt) is approximately 35,640 g/mol.

* Mix gRNA and Cas9 at ratio of 15:1

1. Prepare 5 × Cas9 Cleavage Buffer (5 × CCB, 100 mM HEPES containing 750 mM KCl, 50 mM $MgCl_2$ and 25% glycerol, pH 7.4).
2. Thaw reagents on ice.
3. Dilute gRNA to 3.2 μM (for 64 nM) or 12.8 μM (for 256 nM) or 51.2 μM (for 1024 nM) with DW.
4. Incubate gRNA at 95°C for 2 min → RT for 5 min.
5. Dilute Cas9 to 213.3 nM (for 64 nM) or 853.3 nM (for 256 nM) or 3413.3 nM (for 1024 nM) with 3.3 × CCB.
6. Mix 15 μL of gRNA (or DW for negative control sample) and 15 μL of Cas9 per one reaction, and incubate at 37°C for 10 min.
7. Add 20 μL of gDNA (150 ng/μL) to RNP reaction mixture and mix gently.
8. Incubate at 37°C for 16 h.
9. Add 6.3 μL of RNase A solution to the reaction mixture and mix well.

| | |
|---|---|
| RNase A (10 mg/mL) | 4.4 μL |
| 5 × CCB | 1.4 μL |
| DW | 0.5 μL |
| Total | 6.3 μL |

10. Incubate at 37°C for 20 min.
11. Add 0.5 μL of proteinase K (20 mg/mL).
12. Incubate at 55°C for 20 min.

**F. Purification of RNP digested gDNA**

1. Add 0.1 × volumes of 3 M NaOAC (5.7 μL) and mix gently.
2. Add 1 × volumes of EtOH (125 μL) and invert to mix.
3. Incubate at -20°C for 30 min.
4. Centrifuge at maximum speed and 4°C for 10 min.
5. Wash the pellet with 800 μL of cold 70% EtOH.
6. Discard the EtOH entirely.
7. Resuspend the pellet in 25 μL of DW.

### G. Adapter 1 ligation

| Name | Sequence (5'-3') |
|---|---|
| Adapter1_Fwd | 5'-BioOn-GTTGACATGCTGGATTGAGACTTCCTACACTCTTTCCCTACACGACGCTCTTCCGATCT-3' |
| Adapter1_Rev | 5'-GATCGGAAGAGCGTCGTGTAGGGAAAGAGTGTAGGAAGTCTCAATCCAGCATGTCAAC-3' |

1. Assemble a pair of Adapter 1 oligos according to the following.

| | |
|---|---|
| 2 × annealing buffer (20 mM Tris [pH 7.5], 100 mM NaCl, 2 mM EDTA) | 10 μL |
| Adapter1_Fwd (100 μM) | 1 μL |
| Adapter1_Rev (100 μM) | 1 μL |
| DW | 8 μL |
| Total | 20 μL |

2. Incubate the mixture at 95°C for 5 min, and the cool to RT for ~45 min to form Adapter 1.
3. Assemble the reaction mixture according to the following.

| | |
|---|---|
| 10 × NEB2 | 5μL |
| 10 mM dATP | 5 μL |
| Klenow Exo | 5 μL |
| digested DNA | 25 μL |
| DW | 10 μL |
| Total | 50 μL |

4. Incubate at 37°C for 30 min and store at 4°C until use.
5. Assemble the reaction mixture according to the following.

| | |
|---|---|
| 10 × T4 DNA ligase buffer | 5 μL |
| dA-tailed DNA | 38 μL |
| Assembled adapter 1 | 2 μL |
| T4 DNA Ligase | 5 μL |
| Total | 50 μL |

6. Incubate at 16°C for 16 h**.**

### H. Fragmentation

1. Add 25 μL of SPRIselect reagent to 50 μL of Adapter 1-ligated DNA reaction mixture.
2. Mix by pipetting, and then incubate for 5 min at RT.
3. Pull down the beads using magnetic stand and discard supernatant.
4. Wash the beads with 180 μL of 85% EtOH three times.

5. Discard EtOH entirely, add 50 μL of DW, incubate for 10 min at RT.
6. Pull down the beads using magnetic stand and transfer 45 μL of supernatant to a new tube.
7. Assemble the fragmentation reaction mixture according to the following.

| | |
|---|---|
| Adapter 1 ligated DNA | 40 μL |
| dsFragmentase Buffer v2 (10×) (NEB) | 5 μL |
| dsFragmentase Enzyme (NEBNext dsDNA Fragmentase) | 1.5 μL |
| DW | 3.5 μL |
| Total | 50 μL |

8. Incubate at 37°C for 1 h.
9. Quench reaction by adding 12.5 μL of 0.5 M EDTA.
10. Add 37.5 μL of DW.
11. Add 90 μL of SPRIselect reagent to 100 μL of fragmented DNA sample.
12. Mix by pipetting, and then incubate for 5 min at RT.
13. Pull down the beads using magnetic stand and discard supernatant.
14. Wash the beads with 180 μL of 85% EtOH three times.
15. Discard EtOH entirely, add 50 μL of DW, incubate for 10 min at RT.
16. Pull down the beads using magnetic stand and transfer 48 μL of supernatant to a new tube.

**I. Adapter 2 ligation**

| Name | Sequence (5'-3') |
|---|---|
| Adapter2_N5_Fwd | 5'- PHO-GTCGTATTAGTAGTANNNNNAGATCGGAAGAGCACACGTCTGAACTCC -3' |
| Adapter2_N6_Fwd | 5'- PHO-GTCGTATTAGTAGTANNNNNNAGATCGGAAGAGCACACGTCTGAACTCC -3' |
| Adapter2_N7_Fwd | 5'- PHO-GTCGTATTAGTAGTANNNNNNNAGATCGGAAGAGCACACGTCTGAACTCC -3' |
| Adapter2_Rev | 5'- ACTACTAATACGACT -3' |

1. Assemble the end repair reaction mixture according to the following (NEBNext Ultra End Repair/ dA-tailing module reagents).

| | |
|---|---|
| Fragmented DNA | 27.7 μL |
| End-repair reaction buffer (10×) | 3.3 μL |
| End-repair enzyme mix | 1.5 μL |
| DW | 0.5 μL |
| Total | 33 μL |

2. Incubate at 20°C for 30 min → 65°C for 30 min.
3. During the end-repair reaction, assemble Adapter 2 oligos according to the following.

| | |
|---|---|
| 2 × annealing buffer (20 mM Tris [pH 7.5], 100 mM NaCl, 2 mM EDTA) | 6 μL |
| Adapter2_N5_Fwd (100 μM) | 1 μL |
| Adapter2_N6_Fwd (100 μM) | 1 μL |
| Adapter2_N7_Fwd (100 μM) | 1 μL |
| Adapter2_Rev (100 μM) | 3 μL |
| Total | 12 μL |

4. Incubated at 95°C for 5 min, and then cool slowly to precisely anneal each Adapter 2 oligos.
5. Assemble the Adapter 2 ligation reaction mixture according to the following.

| | |
|---|---|
| Blunt/TA Ligase Master Mix | 7.5 μL |
| Ligase enhancer | 0.5 μL |
| Annealed adapter 2 | 1.25 μL |
| End-repaired DNA | 32.5 μL |
| Total | 41.75 μL |

6. Incubate at 20°C for 30 min→16°C for 16 h.

**J. Affinity purification by streptavidin beads**

1. Prepare 2 × BW buffer (10 mM Tris [pH 7.5], 2 M NaCl, 1 mM EDTA).
2. Wash 25 μL of Dynabeads with 125 μL of 1 × BW buffer per one sample.
3. Incubate at RT for 5 min.
4. Pull down the beads with magnetic stand, discard supernatant.
5. Repeat wash step (step 2-4).
6. Resuspend beads in 41 μL of 2 × BW buffer.
7. Add 41 μL of the resuspended beads to assembled 41.75 μL of adapter 2-ligated DNA mixture.
8. Rotate for 30 min at RT to mix well.
9. Pull down the beads using magnetic stand and discard supernatant.
10. Wash the beads with 200 μL of 1 × BW buffer two times.
11. Wash the beads by 200 μL of 10 mM Tris-HCl (pH 8.5) two times.
12. Resuspend the beads in 20 μL of 10 mM Tris-HCl.

**K. Recovery PCR**

| Oligo name | Sequence (5'-3') |
|---|---|
| Recovery_PCR_Fwd | 5'- GGAGTTCAGACGTGTGCTC -3' |
| Recovery_PCR_Rev | 5'- GTTGACATGCTGGATTGAGACTTC -3' |

1. Assemble the recovery PCR reaction mixture according to the following.

| | |
|---|---|
| 2 × Phusion Master Mix | 25 µL |
| 10 µM Recovery_PCR_Fwd | 2.5 µL |
| 10 µM Recovery_PCR_Rev | 2.5 µL |
| Resuspended Beads Mixture | 20 µL |
| Total | 50 µL |

2. Set up a thermal cycler with the following parameters.

| Step | Temperature | Time | Cycle |
|---|---|---|---|
| 1 | 98°C | 30 sec | 1 |
| 2 | 98°C | 10 sec | 12 |
| | 61°C | 30 sec | |
| | 72°C | 2 min | |
| 3 | 72°C | 2 min | 1 |
| 4 | 4°C | ∞ | 1 |

3. Transfer 30 µL of PCR product using magnetic stand to a new tube.

**L. Indexing PCR**

1. Dilute 3 µL of the recovery PCR product to 148.5 µL of DW.
2. Assemble indexing PCR reaction mixture according to the following.

| | |
|---|---|
| 2 × Phusion Master Mix | 20 µL |
| 5 µM Index primer Forward | 4 µL |
| 5 µM Index primer Reverse | 4 µL |
| Diluted recovery PCR product | 12 µL |
| Total | 40 µL |

3. Set up a thermal cycler with the following parameters.

| Step | Temperature | Time | Cycle |
|---|---|---|---|
| 1 | 98°C | 30 sec | 1 |
| 2 | 98°C | 10 sec | 12 |
| | 61°C | 30 sec | |
| | 72°C | 2 min | |
| 3 | 72°C | 2 min | 1 |
| 4 | 4°C | ∞ | 1 |

**M. Purification of the DNA library**

0.7 × left-side SPRIselect

1. Add 0.7 × volumes of SPRIselect reagent (28 μL) to 40 μL of PCR product.
2. Mix by pipetting, and then incubate for 5 min at RT.
3. Pull down the beads using magnetic stand and discard supernatant.
4. Wash the beads with 180 μL of 85% EtOH three times.
5. Discard EtOH entirely, add 1 × volumes of DW (40 μL), incubate for 10 min at RT.
6. Pull down the beads using magnetic stand and transfer 38 μL of supernatant to a new tube.

0.5 × right-side SPRIselect (to remove >1000 bp large fragments)

1. Add 0.5 × volumes of SPRIselect (19 μL) to 38 μL of sample.
2. Mix by pipetting, and then incubate 5 min at RT.
3. Pull down the beads using magnetic stand, and transfer 55 μL of supernatant to a new tube.
4. Add 1.3 × volumes of SPRIselect (71.5 μL).
5. Mix by pipetting, and then incubate for 5 min at RT.
6. Pull down the beads using magnetic stand and discard supernatant.
7. Wash the beads with 180 μL of 85% EtOH three times.
8. Discard EtOH entirely, add 38 μL of DW and incubate for 10 min at RT.
9. Pull down the beads using magnetic stand and transfer 36 μL of supernatant to a new tube.

0.7 × left-side SPRIselect (to remove residual adapters, <200 bp short fragments)

1. Add 0.7 × volumes of SPRIselect (25 μL) to previous 36 μl of sample.
2. Mix by pipetting, and then incubate for 5 min at RT.
3. Pull down the beads using magnetic stand and discard supernatant.
4. Wash the beads with 180 μL of 85% EtOH three times.
5. Discard EtOH entirely, add 25 μL of DW, and incubate for 10 min at RT.
6. Pull down the beads using magnetic stand and transfer 23 μL of supernatant to a new tube.

**N. Quantification and qualification of the DNA library for sequencing**

Measure the concentration of the library with Qubit HS, and check quality of DNA library with Agilent 2100 Bioanalyzer and High Sensitivity DNA chip.

1. Quantify the amount of each DNA libraries using Qubit HS (x ng/μL).
2. Check the average size of each DNA libraries with Bioanalyzer (y bp).
3. Calculate the concentration (nM) = x (ng/μL) / [y (bp) × 660 (g/mol)] × $10^6$

**O. Sequencing with Illumina MiSeq**

Post-Run Wash before sequencing

1. Boot or reboot the MiSeq instrument, opening the home screen.
2. Prepare 0.5% Tween 20 before use.
3. Push the [Perform Wash] button.
4. Select [Post-Run Wash].
5. Attach a used flow cell on the instrument, and then select [Next].
6. Set the wash cartridge containing 6 mL of 0.5% Tween 20 per well and the 500 mL bottle containing >350 mL of 0.5% Tween 20, respectively.
7. Start wash by pushing [Next] (for 20~30 min).
8. Discard 0.5% Tween 20 in the cartridge and wash bottle.
9. Wash the cartridge with laboratory grade water.
10. Repeat the same wash program (step 3~7).
11. Leave the cartridge and bottle in the instrument after a second wash.

Setting up the sample sheet

1. Create the sample sheet with the illumina Experiment Manager and the following parameters. (http://jp.support.illumina.com/sequencing/sequencing_software/experiment_manager/downloads.html?langsel=/jp/)

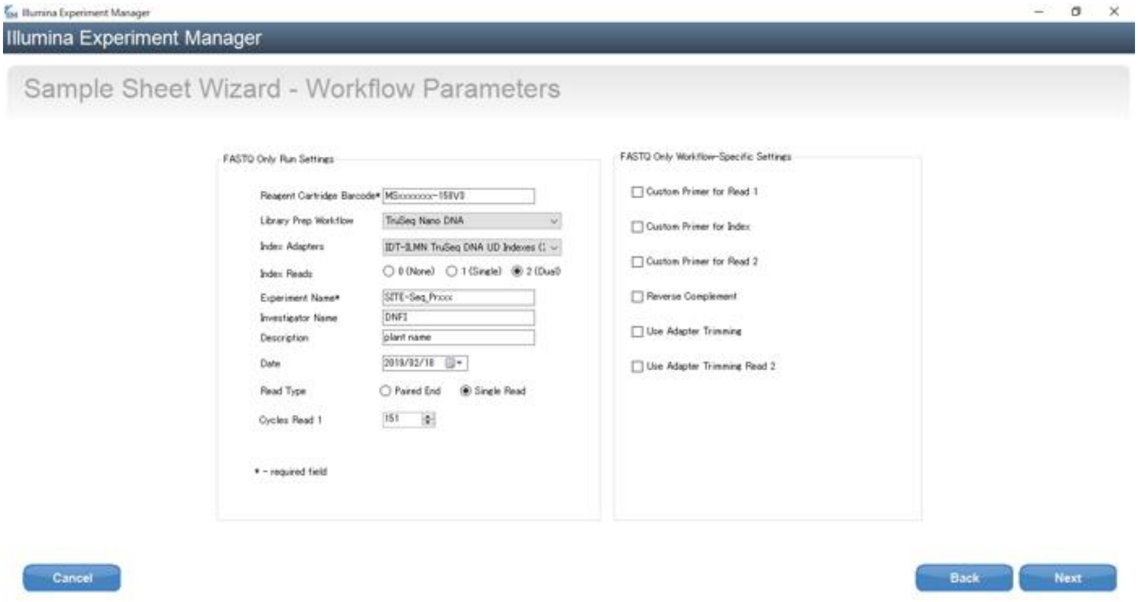


2. Save as “MSxxxxxxx-150V3”, which is the barcode number labeled on the cartridge.
3. Copy and paste the sample sheet into the file location where the sequence data will be saved.

Preparing Reagents

1. Thaw the reagent cartridge.
2. Invert it 10 times.
3. Tap the cartridge on the desk to remove air bubbles in reagents.
4. Store the cartridge at 4°C until use.

Preparing DNA libraries for sequencing

1. Reagents for denaturation and dilution for DNA library are as follows:
   HT1 buffer (supplied in MiSeq kit), thawed and pre-chilled at 4°C
   0.2 N NaOH (dilute before use)
   200 mM Tris-HCl, pH 7.0
2. 1 nM DNA library (20 µL) + 0.2 N NaOH (20 µL) at RT.
3. Mix well, and then incubate for 5 min at RT.
4. Add 20 µL of Tris-HCl, and immediately transfer it on ice.
5. Add 940 µL of ice-cold HT1 (total 1 mL of 20 pM DNA library).
6. 20 pM DNA library (300 µL) + HT1 (300 µL) (total 600 µL of 10 pM DNA library).

   *Final concentration of NaOH must be at $< 0.001$ N.
7. Load 600 µL of 10 pM DNA library into the sample loading port of cartridge (port **No. 17**).

Sequencing with MiSeq

*Set up according to step-by-step instructions on the screen.

1. Reboot the system.
2. Push [Run Option] to specify the location for data saving.
3. Return to home screen, and select [SEQUENCE] to set up the run.
4. Push [Next]. *Leave the check box for the BaseSpace clear.
5. Wash well the flow cell with laboratory grade water to remove salts, and then wipe with a lens cleaning paper prewetted with EtOH.

   *Do not use a paper for the ports of the flow cell.
6. Load the flow cell, and confirm that the RFID is accepted (see the left below corner).
7. Close the door, and click [Next].
8. Set PR2 bottle (provided in the MiSeq kit, stored at 4°C).
9. Discard remaining solution in the waste bottle, and then reset the bottle.
10. Ensure that the RFID of PR2 is accepted, and then select [Next] button.
11. Open the door of the chiller, and insert the reagent cartridge.
12. Close the door immediately, check that the RFID is accepted, and then touch [Next].
13. [Review screen] will display the barcode ID and the corresponding ID of sample sheet.

14. Ensure that a name of experiment and the work flow are correct, and push [Next].
15. After [Pre-Run Check] is finished, click [Start Run] (15 h).

Post-Run Wash after sequencing

1. After successful run, click [Next] button to proceed post-run wash.
2. Prepare 1 L of 0.5% Tween 20 before use.
3. Perform a first wash using 0.5% Tween 20.
4. Check the box, push [Next], and then [Done].
5. Perform a second wash using only 0.5% Tween 20 in the same manner.
6. Discard the flow cell, cartridge and bottle from the instrument.

**Amplicon-sequencing Procedure**

**A. Amplify the target sequences**

1. Synthesize 1st PCR primer set according to the following.

| | |
|---|---|
| Amp_Adapter_Fwd | 5'-CACTCTTTCCCTACACGACGCTCTTCCGATCT[Target]-3' |
| Amp_Adapter_Rev | 5'-GGAGTTCAGACGTGTGCTCTTCCGATCT[Target]-3' |

2. Assemble the PCR reaction mixture according to the following.

| | |
|---|---|
| 5 μM Amp_Adapter_Fwd | 2.5 μL |
| 5 μM Amp_Adapter_Rev | 2.5 μL |
| 2 × Phusion master mix | 12.5 μL |
| Approximately 100 ng of gDNA | x μL |
| DW | (7.5 – x) μL |
| Total | 25 μL |

3. Set up a thermal cycler with the following parameters.

| Step | Temperature | Time | Cycle |
|---|---|---|---|
| 1 | 98°C | 30 sec | 1 |
| 2 | 98°C | 10 sec | 25 |
| | 60°C | 30 sec | |
| | 72°C | 30 sec | |
| 3 | 72°C | 5 min | 1 |
| 4 | 4°C | ∞ | 1 |

4. Load 20 μL of the PCR product on 2% E-gel EX Agarose Gel.
5. Purify the DNA fragment from the agarose gel slice using NucleoSpin Gel and PCR Clean-up.
6. Quantify the amount of DNA with Qubit HS.
7. Dilute DNA sample to 100 pg/μL.

**B. Indexing PCR**

1. Synthesize indexing PCR primer according to the following.

| | |
|---|---|
| Index_UDIxxxx_Fwd | AATGATACGGCGACCACCGAGATCTACACNNNNNNNNACACTCTTTCCCTACACGACG |
| Index_UDIxxxx_Rev | CAAGCAGAAGACGGCATACGAGATNNNNNNNNGTGACTGGAGTTCAGACGTGTGCTC |

2. Assemble the indexing PCR reaction mixture according to the following.

| | |
|---|---|
| 5 µM Index-Fwd | 5 µL |
| 5 µM Index-Rev | 5 µL |
| 2 × Phusion master mix | 25 µL |
| Diluted DNA sample (100 pg/µL) | 1 µL |
| DW | 14 µL |
| Total | 50 µL |

3. Set up a thermal cycler with the following parameters.

| Step | Temperature | Time | Cycle |
|---|---|---|---|
| 1 | 98°C | 30 sec | 1 |
| 2 | 98°C | 10 sec | 10 |
| | 64°C | 30 sec | |
| | 72°C | 30 sec | |
| 3 | 72°C | 5 min | 1 |
| 4 | 4°C | ∞ | 1 |

4. Add 1 × volumes of SPRIselect reagent (40 µL) to 40 µL of amplified DNA sample.
5. Mix by pipetting, and then incubate for 5 min at RT.
6. Pull down the beads using magnetic stand and discard supernatant.
7. Wash the beads using 180 µL of 85% EtOH three times.
8. Discard EtOH entirely, add 20 µL of DW, incubate for 5 min at RT.
9. Pull down the beads using magnetic stand and transfer 18 µL of supernatant to a new tube.

**C. DNA library QC**

1. Quantify the amount of each DNA libraries using Qubit HS.
2. Check the average size of each DNA libraries using Bioanalyzer.
3. Calculate concentration (nM) = x (ng/µL) / [y (bp) × 660 (g/mol)] × $10^6$
4. Assemble each DNA libraries diluted to 1 nM with DW (1 nM DNA library).
5. Dilute 1 nM DNA library to 50 pM with 10 mM Tris-HCl (pH 8.5) (50 pM DNA library).

**D. Sequencing with Illumina iSeq 100**

1. Thaw the cartridge before use with 25°C water bath at least 6 h.
2. Invert the cartridge 5 times and tap the cartridge to collect the reagent to bottom.
3. Load 20 µL of 50 pM DNA library to the Library reserve hole.
4. Set the cartridge to the iSeq100 and perform sequencing with machine instruction.
