## Supplementary Script for "Strategy for detecting off-target sites in genome-edited rice"

```
#!/bin/bash

b_time=`date +%H:%M:%S`

tmp=Fq/tmp
c1=0
c2=1

for f in Fq/*.fastq.gz;
		do
		c1=$((c1+1))
done

for f in Fq/*.fastq.gz;
		do g="${f%%.*}"
		g="${g##*/}"
		echo $c2,¥($c1¥),$g,$f
		c2=$((c2+1))

id=$g

date >> SSprocess.log
echo -e "-U Fq/"${id}".fastq.gz -S sam/"${id}".sam" >>SSprocess.log

fastqc Fq/${id}.fastq.gz

if [ -e IRGSP-1.0_genome.1.bt2 ]; then
		echo 'Index files exist. Skip index build.'
else (bowtie2-build -f IRGSP-1.0_genome.fasta IRGSP-1.0_genome)
fi

bowtie2 ¥
		-x IRGSP-1.0_genome -p 8 ¥
		-U Fq/${id}.fastq.gz ¥
		-S sam/${id}.sam >> SSprocess.log
samtools sort ¥
```

```
        -@ 8 ¥
        -m 4G ¥
        -O bam ¥
        -o sam/${id}.bam sam/${id}.sam ¥
        && mv -f sam/${id}.bam sam/${id}_sorted.bam
samtools index sam/${id}_sorted.bam

minread=10
python siteseq.py ¥
        -i sam/${id}_sorted.bam ¥
        -R IRGSP-1.0_genome.fasta ¥
        -p sam/${id}_mread${minread}_p.txt ¥
        -o sam/${id}_mread${minread}_o_whole.txt ¥
        -t ${minread}
cat sam/${id}_mread${minread}_o_whole.txt ¥
        | awk '!a[$2]++{print $0}' ¥
        | awk 'gsub(":","¥t") {print $3,"¥t",$4,"¥t",$4}' ¥
        | awk 'gsub(" ","") {print $0}' ¥
        | sort -k1,1 -k2,2n >sam/${id}_mread${minread}_o.bed
bedtools intersect ¥
        -a transcripts_exon.gff ¥
        -b sam/${id}_mread${minread}_o.bed ¥
        | awk '$3 == "mRNA" {print $0}' >sam/${id}_mread${minread}_o_annotation.txt

        cp $f $tmp
        rm -f $f
        rm -f sam/$g.sam

done
echo START-END=${b_time}-`date +%H:%M:%S`
```
